# Microbiota-derived indole limits *Campylobacter jejuni* colonization by inhibiting respiration and metabolism

**DOI:** 10.64898/2026.04.28.721463

**Authors:** Ritam Sinha, Barsha Bhattarai, Cristina Kraemer Zimpel, Elizabeth Ottosen, Rhiannon M. LeVeque, Pallavi Singh, Victor J. DiRita

## Abstract

How the microaerophilic *Campylobacter jejuni* grows in the inflamed gut is poorly understood, hampering our ability to identify novel targets for controlling infections by this pathogen. Our prior ferret study indicated that *C. jejuni* growth during infection is driven by intestinal inflammation. Without manipulating microbiota or introducing pro-inflammatory genetic lesions (or both), conventional mice are naturally resistant to *C. jejuni* colonization and infection. To test the impact of inflammation *per se* on *C. jejuni* infection, we induced transient intestinal inflammation in mice using short-term dextran sodium sulfate (DSS) treatment. The resulting colitis disrupted colonization resistance, enabling rapid *C. jejuni* growth in the murine colon within three days of infection, accompanied by exacerbated intestinal inflammation. DSS-induced colitis led to enrichment of mucin-degrading bacteria and depletion of taxa producing short-chain fatty acids and indole; metabolomic profiling confirmed a marked reduction in colonic indole levels in both DSS-treated and infected mice. At physiological concentrations, indole inhibited *C. jejuni* growth *in vitro* and resulted in reduced transcript levels from key energy-generating pathways, including nitrate respiration (*napA*), aerobic respiration (*ccoN*), lactate utilization (*lctP*), and the acetate switch (*ackA/ptaA*); consistent with this, mutations in these pathways led to fitness defects in DSS-treated mice, highlighting their importance for *C. jejuni* colonization in the inflamed gut. Moreover, treatment with indole or the indole-producing probiotic *Escherichia coli* Nissle 1917 significantly reduced *C. jejuni colonization in vivo*. Our findings show how C. jejuni establishes and grows in the inflamed gut and demonstrate that microbiota-derived metabolites play a key role in regulating *C. jejuni* pathogenicity.

**Significance:** The mechanisms by which the microaerophilic pathogen *C. jejuni* proliferates in the inflamed intestine remain poorly understood. Using a DSS-induced colitis mouse model, we identify host physiological changes, microbiota alterations, and metabolic factors that promote *C. jejuni* colonization during intestinal inflammation. These findings provide mechanistic insight into why *Campylobacter* species are frequently detected in patients with Inflammatory Bowel Disease (IBD) and how infection can exacerbate intestinal inflammation and worsen disease symptoms. We further identify the microbiota-derived metabolite indole as a potent inhibitor of *C. jejuni* colonization, highlighting a potential metabolite-based therapeutic strategy to combat emerging multidrug-resistant *C. jejuni* infections.

## Introduction

*Campylobacter jejuni*, a Gram-negative foodborne pathogen, is a leading cause of gastroenteritis in humans, primarily transmitted via contaminated food products (1). The emergence of multidrug-resistant strains highlights the urgent need to understand the molecular mechanisms underlying *C. jejuni* pathogenesis (2). A central question in this field is how *C. jejuni* adapts and proliferates in the face of host inflammation.

We demonstrated that the ferret is a physiologically relevant model of *C. jejuni* disease and determined that the pathogen exploits intestinal inflammation to grow in the gut (3). During acute infection in this model, *C. jejuni* triggered pronounced crypt hyperplasia in the colon, consistent with activation of epithelial repair programs. This response was accompanied by goblet cell depletion and a marked increase in undifferentiated, proliferating (Ki67+) epithelial cells. We also observed increased epithelial oxygenation and metabolic remodeling, including elevated colonic luminal L-lactate, which supported bacterial growth (3). Although highly informative, the ferret model is constrained by high cost, limited tractability, and the lack of robust genetic tools to dissect host determinants of *C. jejuni* proliferation.

Mice, in contrast, offer a well-characterized immune system, genetic tractability, and affordability. However, to support *C. jejuni* colonization and growth, conventional mouse models typically require genetic modifications, such as IL-10 deficiency, or manipulation of the microbiota, complicating studies of host-pathogen interactions (4–6). Based on our observations using ferrets to study *C. jejuni* infection, we hypothesized that inducing an intestinal inflammatory environment in mice would permit *C. jejuni* colonization and growth without genetic or antibiotic interventions.

In this study, we applied short-term treatment with dextran sodium sulfate (DSS), a widely used inflammatory agent in murine models of colitis and irritable bowel syndrome, leading to increases in epithelial oxygenation and luminal lactate levels (7–9). Following DSS treatment, mice were orally infected with *C. jejuni*, resulting in successful colonization and growth, demonstrating that inflammation alone is sufficient to facilitate this outcome. We further characterized the DSS-treated mouse model by profiling gut microbiota and metabolites. Notably, DSS treatment reduced levels of indole-producing commensals, leading to a significant decrease in luminal indole. Further analysis revealed that indole directly inhibits *C. jejuni* growth and colonization by disrupting bacterial respiration and metabolic pathways, thereby highlighting a previously unappreciated microbiota-mediated mechanism of colonization resistance. Together, these findings establish a physiologically relevant and genetically tractable mouse model of *C. jejuni* colonization driven by inflammation. This model provides a robust platform to dissect host, microbial, and metabolic factors that regulate pathogen growth and offers insights into how *C. jejuni* exploits inflammatory niches to overcome microbiota-mediated resistance.

## Results

### Intestinal inflammation promotes robust *C. jejuni* growth in the murine colon

Intestinal inflammation promotes the outgrowth of several facultative enteric bacteria, including *Salmonella* Typhimurium, *E. coli*, and *Citrobacter rodentium*, in the murine gut (10–12). However, whether inflammation similarly enhances colonization by the microaerophilic pathogen *C. jejuni* remains poorly understood and our work with a ferret model led us to consider this possibility.

We performed an infection experiment using four groups of female mice (n = 5 per group). Two groups were treated with 3% DSS in drinking water for three days to induce intestinal inflammation, while the remaining two groups received normal drinking water as controls. On the following day, one DSS-treated group and one control group were orally infected with *C. jejuni* WT 11168 strain (10⁸ CFU/mL). After infection, DSS was withdrawn and all mice were returned to normal drinking water **(Fig. 1A)**.

**Figure 1.**
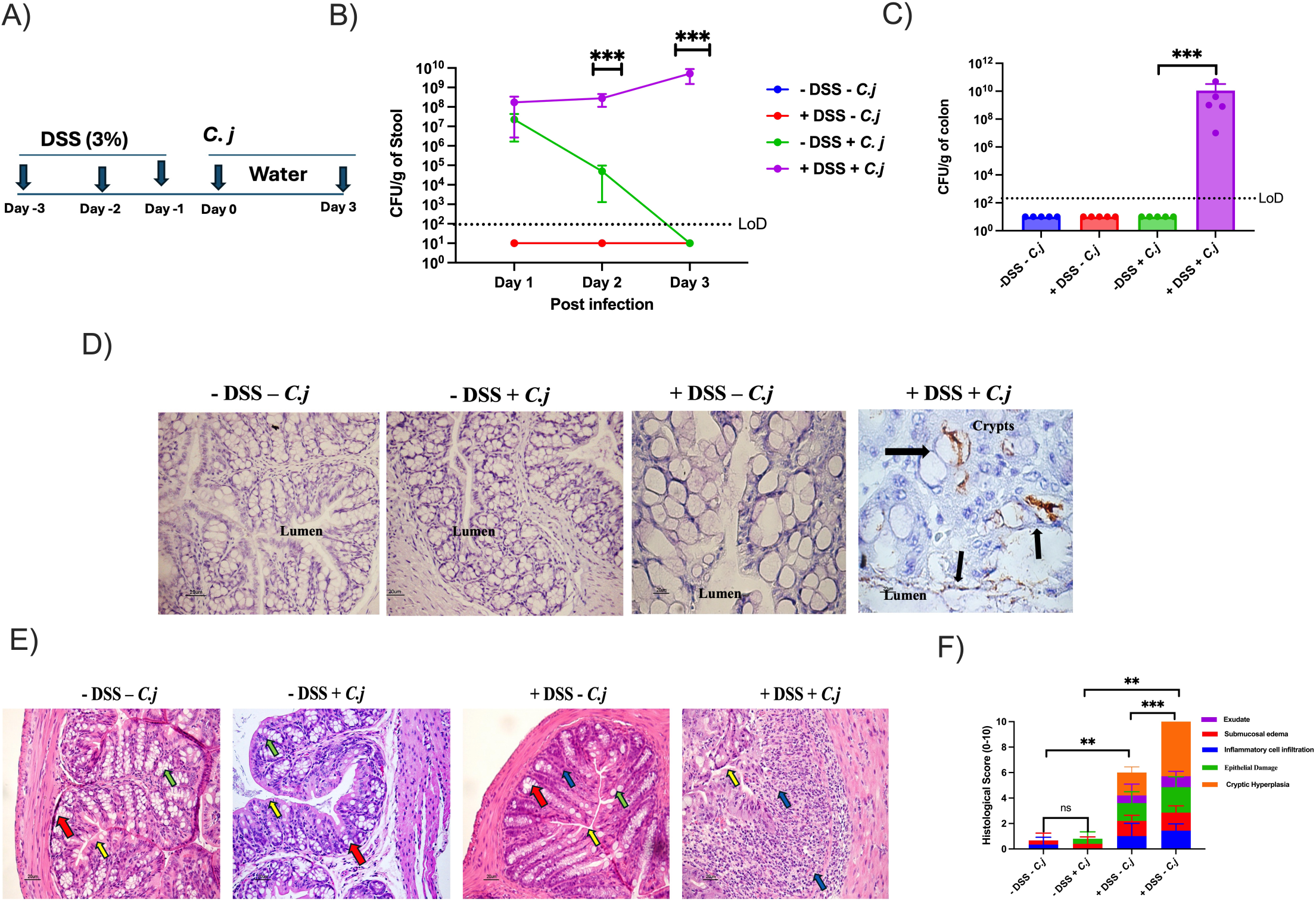
Intestinal inflammation promotes *C. jejuni* growth in the mouse gut. (A) Schematic diagram of the experimental design for the DSS-induced colitis and *C. jejuni* infection mouse model. (B) *C. jejuni* 11168 WT bacterial loads in stool samples were quantified by colony-forming unit (CFU) counts on days 1, 2, and 3 post-infection following oral inoculation with 10^8^ CFU/mL (n = 5). (C) *C. jejuni* 11168 WT loads in colon tissues were determined by CFU counts on day 3 post-infection (n = 5). (D) Immunohistochemistry (IHC) using a *C. jejuni*-specific antibody was performed to determine bacterial localization in colon tissue on day 3 post-infection. Black arrows indicate the presence of *C. jejuni* in infected colonic tissue. Representative images are shown from two independent experiments. (E) Colonic inflammatory responses were evaluated by histological analysis and histological scoring on day 3 post-infection. Representative hematoxylin and eosin (H&E)–stained sections are shown. Yellow arrows indicate epithelial cells, red arrows indicate intestinal crypts, green arrows indicate goblet cells, and blue arrows indicate infiltrating leukocytes in the lamina propria. Data are presented as mean ± SE. Statistical analysis was performed using one-way ANOVA. *\*P < 0.05; **P < 0.01; ***P < 0.001; ****P < 0.000*.

At day one post-infection, fecal loads of *C. jejuni* were approximately 10⁸ CFU/g in both DSS-treated and untreated infected groups. However, by day three, fecal *C. jejuni* levels gradually declined in DSS-untreated infected mice, whereas DSS-treated infected mice showed an increase to approximately 10⁹-10^10^ CFU/g **(Fig. 1B)**. Because peak colonization in human infection is typically observed two-to-three days post-infection in the large intestine, we next quantified *C. jejuni* loads in the colon at day three. We detected approximately 10⁹–10¹⁰ CFU *C. jejuni*/g of colon tissue in DSS-treated infected mice **(Fig. 1C).** In contrast, *C. jejuni* was undetectable or below the limit of detection in DSS-untreated infected mice, as well as in both uninfected control groups. These results indicate that DSS-induced inflammation promotes *C. jejuni* growth in the mouse colon by day three post-infection. The highest colonization was observed in the colon and cecum (∼10⁹ CFU/g tissue), whereas the small intestine showed lower levels (∼10⁶ CFU/g tissue) at day three post-infection **(FigS1A)**. Because *C. jejuni* levels were highest in the colon, we examined its localization within colonic tissue. Immunohistochemistry with an anti-*Campylobacter* antibody revealed sparse clusters of *C. jejuni* in the colonic lumen, closely associated with the colonic epithelium and detectable within crypts. This localization pattern resembles what has been reported during both human and ferret infection **(Fig. 1D)**(3, 13). *C. jejuni* colonized across the colon, with no significant difference between proximal and distal segments (∼10⁷ CFU/g tissue) **(FigS1B)**. We also examined whether colonization was sex-dependent by comparing male and female mice and found no significant difference between sexes **(FigS1C)**.

We evaluated the inflammatory response following DSS treatment and *C. jejuni* infection. H&E staining and histopathological scoring showed a moderate inflammatory response from DSS treatment alone, characterized by disruption of the epithelial cell layer, leukocyte infiltration into the superficial lamina propria, cryptic edema and cryptic hyperplasia in colonic tissue compared to the two control groups. Infection with *C. jejuni* in DSS-treated mice triggered a more severe inflammatory response, with significantly higher pathological scores and more pronounced tissue damage compared to DSS treatment alone **(Fig. 1D)**.

To further characterize the inflammatory environment, we measured colonic mRNA levels of key cytokines using RT-PCR. are Established proinflammatory mediators in colitis are IL-6, TNF-α, and KC: IL-6 promotes immune cell activation and sustains chronic inflammation, TNF-α drives epithelial damage and leukocyte recruitment, and KC (CXCL1) recruits neutrophils that contribute to mucosal injury(14, 15). In contrast, IL-10 is a potent anti-inflammatory cytokine that limits excessive immune responses in the gut by suppressing proinflammatory signaling, preserving barrier function, and maintaining immune tolerance to microbial antigens; loss of IL-10 leads to more severe colitis in animal models(16). Transcript levels were compared among DSS-treated uninfected mice, DSS-treated infected mice, and untreated uninfected controls. As expected, DSS treatment alone induced a strong inflammatory response, with ∼five-to-15-fold increased transcription of IL-6, TNF-α, and KC, along with an ∼five-fold reduction in IL-10 expression compared to untreated uninfected mice **(Fig. 2A)**. *C. jejuni* infection in DSS-treated mice amplified this response. Relative to DSS-treated uninfected mice, infected mice showed elevated expression of all three pro-inflammatory cytokines, while IL-10 expression was further reduced **(Fig. 2A)**.

**Figure 2.**
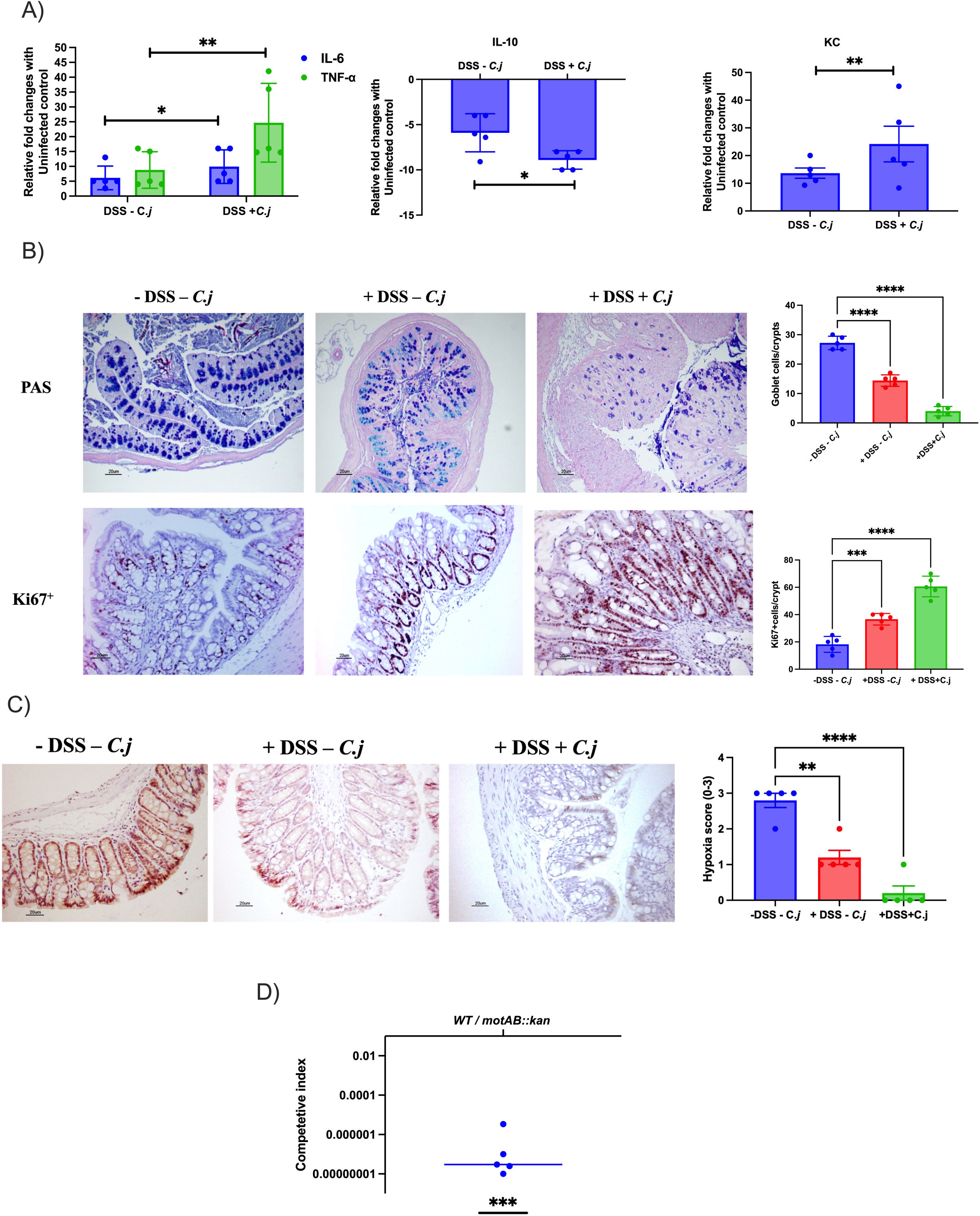
Inflammatory and pathophysiological changes during *C. jejuni* infection in DSS-treated mice. (A) Relative fold changes in proinflammatory cytokine genes (*IL-6, TNF-α,* and *KC*) and the anti-inflammatory cytokine gene (*IL-10*) were determined by qRT-PCR analysis of colon tissues from DSS-treated uninfected and *C. jejuni*-infected mice, compared with the PBS control group (n = 5). Gene expression changes were calculated using the 2^−ΔΔ*CT*^ method. Statistical analysis was performed using one-way ANOVA. *P* < 0.05; P < 0.01; *P* < 0.001; P < 0.0001. (B) Representative images of Alcian blue staining showing goblet cells in colonic tissue and quantification of goblet cells per crypt on day 3 post-infection (n = 5) and representative images showing localization of Ki67⁺ proliferating cells and quantification of Ki67⁺ cells per crypt in colonic tissue on day 3 post-infection (n = 5). Statistical analysis was performed using one-way ANOVA. *P* < 0.05; P < 0.01; *P* < 0.001; P < 0.0001. (C) Hypoxia in colonic tissue was assessed using pimonidazole staining. Binding of pimonidazole (hypoxia probe) was detected by immunohistochemistry (IHC) using an anti-pimonidazole antibody in colon sections collected on day 3 post-infection. Hypoxia scores were determined by blinded scoring of staining intensity in colon sections. Statistical analysis was performed using one-way ANOVA. *P* < 0.05; P < 0.01; *P* < 0.001; P < 0.0001. (D) Competitive infection analysis between the WT strain and the *motAB::kan* mutant. DSS-treated mice were infected with an equal ratio (1:1) of WT and *motAB::kan* mutant strains. On day 3 post-infection, the ratio of mutant to WT bacteria was determined and expressed as the competitive index (CI). Statistical analysis was performed using a one-sample t-test against a hypothetical value of 1 (n = 5 per group). All data are presented as mean ± SD. Statistical analyses were performed using an unpaired two-tailed t-test unless otherwise indicated. *\*P < 0.05; **P < 0.01; ***P < 0.001; ****P < 0.000*.

Infection with *C. jejuni* in the ferret induces significant epithelial differentiation and proliferation(3). To determine whether DSS-induced inflammation and *C. jejuni* colonization results in similar hyperplasia in the mouse, we quantified goblet cells and Ki67⁺ proliferative cells in colonic tissue at day three post-infection. DSS treatment alone significantly reduced the number of Alcian blue–positive goblet cells per crypt, with an approximately two-fold reduction relative to untreated uninfected controls **(Fig. 2B)**. This effect was enhanced after *C. jejuni* infection, with the DSS-treated, infected mice exhibiting approximately three-fold reduction in goblet cells per crypt compared with controls, indicating a significant loss of mucus-producing, terminally differentiated cells during infection **(Fig. 2B)**.

We next assessed epithelial proliferation using Ki67 immunohistochemistry. DSS treatment alone increased by almost two-fold the number of Ki67⁺ cells per crypt compared to untreated uninfected mice, consistent with activation of epithelial repair. The Ki67⁺ cell numbers were further increased to approximately three-fold after *C. jejuni* infection of DSS-treated mice, demonstrating that infection amplifies crypt hyperplasia and expansion of proliferating epithelial cells **(Fig. 2B).** This phenotype closely mirrors the epithelial remodeling response we previously observed during *C. jejuni* infection in ferrets (3).

A major consequence of epithelial differentiation state is its impact on oxygen consumption and oxygen availability at the mucosal surface(10). Mature colonocytes metabolize microbially derived butyrate through β-oxidation, a process that consumes oxygen and contributes to maintenance of physiological hypoxia in the intestinal lumen(10). In contrast, undifferentiated proliferating epithelial cells, enriched at the base of crypts, lack robust β-oxidation and rely on aerobic glycolysis, producing lactate from glucose. Because these proliferating cells consume less oxygen, crypt bases are relatively more oxygenated than the upper crypt regions, and expansion of undifferentiated epithelial populations is predicted to increase epithelial oxygenation overall(10).

Because DSS treatment and *C. jejuni* infection both increased Ki67⁺ proliferating cells and reduced terminally differentiated goblet cells, we hypothesized that these conditions disrupt physiological epithelial hypoxia and lead to elevated epithelial oxygenation during acute infection. To test this, we used pimonidazole staining, a well-established *in vivo* hypoxia probe. Under hypoxic conditions, pimonidazole is reduced to reactive intermediates that form stable adducts with proteins and DNA, allowing hypoxic regions to be detected by immunohistochemistry using anti-pimonidazole antibodies.

Mice were injected intraperitoneally with pimonidazole one hour before euthanasia. Mice without either DSS treatment or *C. jejuni* infection exhibited strong pimonidazole retention in colonic epithelial cells, consistent with maintenance of a hypoxic epithelial environment under steady-state conditions. Uninfected, DSS-treated mice showed reduced pimonidazole staining, indicating partial loss of epithelial hypoxia during DSS-induced colitis. *C. jejuni*-infected, DSS-treated mice showed near-complete loss of pimonidazole signal, consistent with severe disruption of epithelial hypoxia and increased epithelial oxygenation **(Fig. 2C)**. These results demonstrate that DSS-induced colitis partially disrupts epithelial hypoxia, and that *C. jejuni* infection further exacerbates this loss, consistent with our previous observations linking *C. jejuni* infection to increased epithelial oxygenation (3).

We observed that *C. jejuni* can reproducibly colonize and grow in the murine intestinal tract after DSS treatment. As a proof of principle to determine whether this DSS inflammation model can be used to identify bacterial fitness factors, we performed an *in vivo* competition assay using the wild-type strain and a mutant deficient for the genes (*motAB*) encoding the established fitness determinant *motAB*, which regulates motility and is critical for colonization in other models(17). Following DSS treatment, mice were orally inoculated with a 1:! mixture of wild-type *C. jejuni* 11168 and a *motAB::kan* derivative at a total dose of 10⁸ CFU. Colonization was assessed in the colon at day three post-infection. In this competitive setting, the *motAB::kan* mutant exhibited a dramatic colonization defect compared with the wild-type strain, demonstrating that motility provides a strong fitness advantage in the inflamed gut environment **(Fig. 2D)**. These results validate the DSS-treated mouse model as a robust system for identifying *C. jejuni* fitness factors required for colonization during intestinal inflammation.

Overall, our findings demonstrate that DSS treatment disrupts intestinal homeostasis by inducing epithelial injury and mucosal inflammation, overcoming colonization resistance and creating a permissive niche for *C. jejuni* colonization and growth. This model shows that transient inflammatory perturbation alone is sufficient to render conventional mice susceptible to robust *C. jejuni* growth without the need for prior antibiotic treatment or genetic manipulation of the host. We also observed that *C. jejuni* further exacerbates intestinal inflammation following DSS exposure, amplifying mucosal pathogenicity and pro-inflammatory responses.

### Composition of the DSS-treated microbiota and its associated metabolites promote *C. jejuni* **colonization**

The resident microbiota of conventional mice is a key determinant of colonization resistance against *C. jejuni*(18), which readily colonizes germ-free and antibiotic-treated mice(18, 19). Given this established barrier function of the microbiota, we investigated whether DSS-induced inflammation alters the intestinal microbial community in a manner that could reduce colonization resistance and facilitate *C. jejuni* growth. To assess whether DSS treatment changes the overall microbial community structure and whether *C. jejuni* colonization further modifies this environment, we performed 16S rRNA gene sequencing of the V3-V4 region from colonic contents collected at day three post-infection. Three experimental groups were analyzed: DSS-treated *C. jejuni*–infected mice (n = 5), DSS-treated uninfected mice (n = 5), and DSS-untreated uninfected controls (n = 3).

We first evaluated within-sample microbial diversity using alpha diversity metrics, including observed richness, Shannon diversity index, and Chao1. Across all three measures, we observed no significant changes **(SFig.2 A)**, indicating that DSS treatment and *C. jejuni* colonization do not substantially alter overall microbial richness or evenness at day three. suggesting that the ability of DSS-treated mice to support *C. jejuni* growth is not driven by a global loss of microbial diversity.

Analysis of beta diversity using Bray–Curtis dissimilarity, which accounts for relative taxon abundance, and Jaccard distance, which is based on presence or absence of taxons, revealed significant differences in microbial community composition among experimental groups. Principal coordinate analysis (PCoA) derived from both metrics demonstrated clear separation of DSS-treated mice, whether infected or uninfected, from untreated controls **(Fig. 3A,B & SFig.2 B,C)**. This indicates that DSS treatment is the primary driver of microbiota restructuring, whereas *C. jejuni* colonization does not significantly alter overall community composition beyond DSS-induced changes.

**Figure 3.**
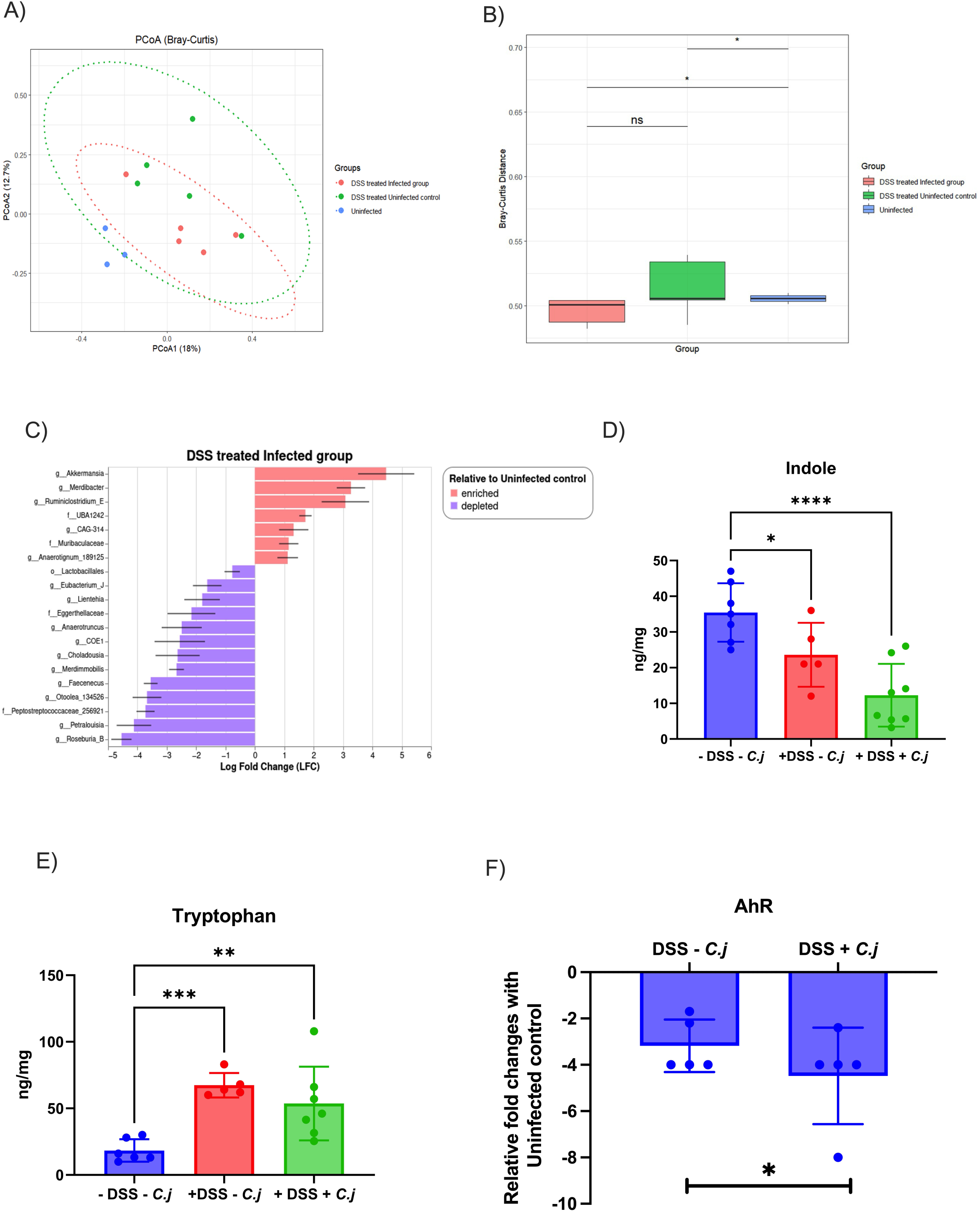
Microbial and metabolic alterations during *C. jejuni* infection in DSS-treated mice. Microbial composition of colon contents from control, DSS-treated uninfected, and *C. jejuni*-infected mice was analyzed by 16S rRNA gene sequencing. (A) Principal coordinate analysis (PCoA) based on Bray–Curtis dissimilarity showing distinct clustering of uninfected controls (n=3) compared with DSS-treated uninfected (n=5) and infected groups (n=5). (B) Boxplots of Bray–Curtis distances demonstrating significant differences in microbial community structure between uninfected controls and DSS-treated uninfected and infected groups. Statistical significance was determined by PERMANOVA (999 permutations) with pairwise post hoc testing. (C) Differential abundance analysis of bacterial genera in DSS-treated groups relative to uninfected controls. Log fold change (LFC) analysis highlights genera significantly enriched (red bars) or depleted (purple bars) in DSS-treated infected groups compared with controls. (D) Concentrations of indole and tryptophan in colon contents from control (n=4), DSS-treated uninfected(n=5), and infected mice (n=6) were determined by mass spectrometry on day 3 post-infection. Statistical analysis was performed using one-way ANOVA. (D) Relative fold changes in AhR gene expression were determined by qRT-PCR analysis of colon tissues from DSS-treated uninfected and *C. jejuni*-infected mice compared with PBS controls (n = 5). Gene expression changes were calculated using the 2^−ΔΔ*CT*^ method. Data are presented as mean ± SD. Statistical analysis was performed using one-way ANOVA unless otherwise indicated. *\*P < 0.05; **P < 0.01; ***P < 0.001; ****P < 0.000*.

Differential abundance analysis (ANCOM-BC) using untreated controls as the reference revealed pronounced microbiota changes following DSS treatment, again indicating that DSS-induced dysbiosis is the primary driver of microbial remodeling in this model. In DSS-treated mice, *Akkermansia* and *Meridibacter* were significantly enriched, likely accounting for increased mucin degradation and altered gut barrier interactions (**SFig.2 D**)(20, 21). DSS-treated *C. jejuni* infected groups showed a similar pattern, with enrichment of *Akkermansia* and *Meridibacter* and deeper depletion of SCFA producers. Although most of the changes we identified were correlated to DSS treatment, we note that *Ruminiclostridium_E* and members of the *Muribaculaceae* family were significantly enriched only in the DSS-treated infection group, suggesting infection-specific taxonomic shifts (**Fig. 3C)**(**22, 23**).

The DSS-treated groups were depleted for a broad set of obligate anaerobic commensals, including *Anaerotruncus*, *Roseburia_B*, *Lachnoclostridium_B*, multiple *Eubacterium* clades, *Peptoclostridium_A*, *Faecenecus*, and members of the *Oscillospiraceae* family (all log-fold change < −2, with false discovery rates of < 0.05) (**SFig.2 D &** (**Fig. 3C)**). Many of these taxa are major producers of short-chain fatty acids (SCFA), particularly butyrate, essential for maintaining epithelial barrier integrity, mucus production, and immunoregulatory signaling(23, 24).

In addition to SCFA production, several Clostridia-associated taxa, including *Anaerotruncus*, *Oscillibacter*, *Eubacterium*, and *Lachnoclostridium*, participate in amino acid fermentation pathways that generate indole and related metabolites (**Fig. 3C)** (**25, 26**). Certain anaerobes, including *Peptostreptococcus* species, convert dietary tryptophan into bioactive indole derivatives such as indoleacrylic acid (IA) and indole-3-propionic acid (IPA) (27). The positive association between *Eubacterium* abundance and levels of indole metabolites further underscores the contribution of these taxa to microbial tryptophan metabolism(25). Indole and its derivatives support gut homeostasis by strengthening epithelial barrier function and regulating mucosal immunity as ligands of the Aryl hydrocarbon Receptor (AhR), which mediates anti-inflammatory signaling (28).

We conclude that DSS treatment induced a dysbiotic microbiota marked by expanding populations of mucin-degrading organisms and depletion of SCFA- and indole-producing obligate anaerobes, thereby creating a gut environment permissive for *C. jejuni* colonization, with reduced microbial support for intestinal homeostasis, while infection primarily amplified changes in select taxa rather than generating a distinct community configuration.

Because several indole-producing bacterial taxa were significantly reduced by DSS in both infected and uninfected groups, we hypothesized that luminal indole levels would be correspondingly diminished in these mice. To test this, we quantified indole and L-tryptophan concentrations in colonic samples by LC/MS mass spectrometry, as gut microbiota convert L-tryptophan into indole and related derivatives through tryptophan metabolism. Consistent with the loss of indole-producing taxa, indole levels were significantly decreased in both DSS-treated uninfected and infected groups compared with untreated controls, whereas tryptophan levels showed a relative accumulation, suggesting impaired microbial conversion of tryptophan to indole (**Fig. 3D,E)**.

The aryl hydrocarbon receptor (AhR) is a key transcription factor that regulates epithelial barrier integrity, IL-22, IL-10 expression, and mucosal immune homeostasis (29–31). Given reduced levels of the AhR ligand indole, we assessed AhR transcriptional expression in colonic tissues by RT PCR. *Ahr* mRNA levels were significantly reduced in both DSS-treated groups, with an approximately four-fold reduction in DSS-treated uninfected mice and an approximately eight-fold reduction in DSS-treated infected mice compared with untreated controls (**Fig. 3F)**.

Overall, DSS treatment reshaped the gut microbiota, resulting in depletion of SCFA- and indole-producing taxa, consistent with reduced butyrate availability and increased epithelial oxygenation in both DSS-treated groups. Indole levels were significantly decreased, accompanied by reduced expression of AhR and IL-10. Because AhR activation promotes IL-10–mediated mucosal tolerance(31), and IL-10 impacts *C. jejuni* colonization, diminished indole availability may facilitate *C. jejuni* pathogenicity through impaired immune regulation. It may also result in loss of direct antimicrobial pressure, as indole exhibits antibacterial and virulence activity against multiple enteric pathogens (32–34).

### Microbiota derived indole inhibits *C. jejuni* growth *in vitro* and *in vivo*

Indole is microbiota-derived tryptophan metabolite that accumulates in the colon at physiological concentrations (∼0.25–1 mM) (32). At these levels, it supports intestinal homeostasis by strengthening epithelial barrier function as a ligand of AhR, regulating mucosal immunity, and exerting antimicrobial activity by disrupting membrane potential and respiration (32, 35). Given that indole levels were markedly reduced following DSS treatment and during *C. jejuni* infection, we investigated whether indole directly influences *C. jejuni* growth, as its effect on this pathogen remains poorly defined. For this, wild-type *C. jejuni* was cultured in Mueller–Hinton broth supplemented with physiologically relevant concentrations of indole (0.25–1 mM), with DMSO serving as the vehicle control, and bacterial growth was assessed by CFU enumeration over a 30-hour time course.

There was little effect on *C. jejuni* growth in 0.25 mM indole, while significant growth reduction was detected as early as six hours in 1 mM indole, and by nine hours in 0.5 mM indole (**Fig. 4A)**.. In these latter two concentrations, bacterial viability progressively declined so that by 30 hour of cultivation in 1 mM indole *C. jejuni* exhibited near-total loss of viability, an observation that was not limited to strain 11168 but was also observed in three additional *C. jejuni* isolates from different sources tested under the same conditions **(SFig. 3A)**. These findings demonstrate that indole at physiological concentrations inhibits *C. jejuni* growth *in vitro*.

**Figure 4.**
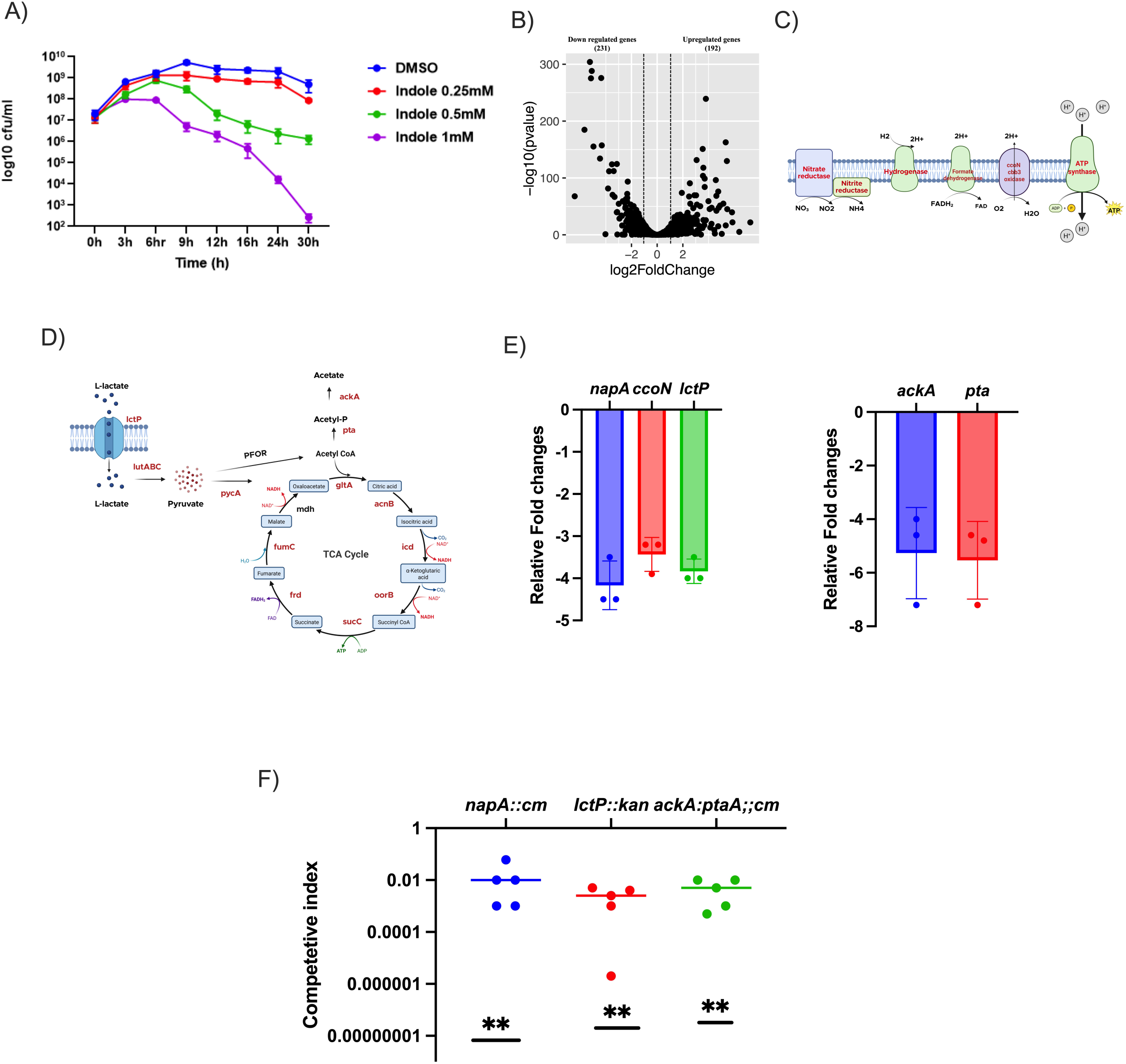
Indole inhibits *C. jejuni* growth by suppressing respiratory and metabolic pathways. (A) *C. jejuni* growth was determined by colony-forming unit (CFU) counts at different time intervals up to 30 h in the presence or absence of indole at physiological gut concentrations (0.25–1 mM) under in vitro conditions. (B) Transcriptomic analysis of *C. jejuni* gene expression after 9 h of growth in the presence of indole (0.5 mM) or vehicle control (DMSO) was performed by RNA-seq (n = 3). A volcano plot shows significantly upregulated and downregulated genes (>2-fold change) in response to indole treatment. (C–D) Representative pathway diagrams highlighting significantly downregulated respiratory and metabolic genes of C. jejuni (shown in red) in response to indole treatment. (E) Quantitative RT-PCR analysis of genes involved in respiration and metabolism, including nitrate reductase (*napA*), cytochrome c oxidase (*ccoN*), lactate transporter (*lctP)*, and acetogenesis pathway genes (*ackA* and *pta*), following 9 h growth in the presence or absence of indole (0.5 mM) under anaerobic and microaerobic conditions. Gene expression changes were calculated using the method. (F) Competitive infection analysis between the WT strain and metabolic mutants (*napA::cm*, *lctP::kan*, and *ackA::pta::cm*). DSS-treated mice were infected with an equal ratio (1:1) of WT and mutant strains. On day 3 post-infection, the ratio of mutant to WT bacteria was determined and expressed as the competitive index (CI). Statistical analysis was performed using a one-sample t-test against a hypothetical value of 1 (n = 5 per group).Data are presented as mean ± SD from three independent experiments unless otherwise indicated. Statistical analysis was performed using one-way ANOVA or an unpaired two-tailed t-test as appropriate. *P < 0.05; **P < 0.01; ***P < 0.001; ****P < 0.000.

To investigate the molecular mechanisms underlying indole-mediated growth inhibition, we performed transcriptomic analysis using RNA sequencing (RNA-seq). Based on our growth kinetics data, we selected the nine-hour time point for analysis, as a significant growth defect was first observed at this stage in the presence of 0.5 mM indole, without loss of viability (**Fig. 4A)**. Wild-type *C. jejuni* was cultured in Mueller–Hinton broth with 0.5 mM indole or DMSO control, and total RNA was isolated at nine hours for transcriptomic profiling. We identified 231 transcripts with lower abundance and 192 genes with greater abundance in the presence of indole (log 2>1, P<0.05) (**Fig. 4B)**. Several respiratory regulatory gene mRNAs were reduced by more than twofold, including those involved in aerobic respiration (e.g., *ccoNPOQ*, encoding a terminal oxidase), nitrate respiration (*napAGHBLD*), hydrogenase (*hydAB*), and formate hydrogenase (*fdhA* and *fdhB*) (**Fig. 4C, Supplemental table 1**).

*C. jejuni* lacks a complete glycolytic pathway but possesses a complete citric acid (TCA) cycle, which serves as its primary metabolic pathway(36). Our data show that transcripts associated with TCA cycle intermediates (*pycA, gltA, acnB, oorC, fumC, mqo, sucC, frdC, and frdB*) were significantly less abundant (> 2 fold) in the presence of indole (**Fig. 4D, Supplemental table 1)**.

The acetate switch in *C. jejuni*, which converts acetyl-CoA to acetate and represents a major ATP-generating pathway, was also affected: transcripts from both *pta* and *ackA* were less abundant by more than two- to four-fold in the presence of indole(37). In addition, those of the lactate transporter *lctP* and the *lutABC* lactate oxidoreductase complex, which converts lactate to pyruvate, were reduced by three- to six-fold (**Fig. 4D, Supplemental table 1)**. Lactate serves as a major carbon source supporting *C. jejuni* growth during inflammation in the ferret (3). To validate the RNA-seq results, we measured *napA*, *ccoN*, *ackA*, *pta*, and *lctP* expression under the same conditions by RT-PCR. We observed an approximately sixfold decrease in *napA*, a fourfold reduction in *ccoN,* and nearly eightfold decreases in both *ackA* and *pta* in the presence of indole, confirming the RNASeq findings **(Fig. 4E)**. Collectively, these data indicate that indole exposure induces a coordinated reduction of respiratory capacity and central metabolic pathways, providing a mechanistic basis for the observed growth inhibition of *C. jejuni*.

To validate the transcriptomic findings further, we assessed bacterial respiration using an assay that measures reduction of 2,3,5-triphenyl-tetrazolium chloride (TTC). Wild-type *C. jejuni* was grown for nine hours in the presence of indole (0.5mM) or DMSO. Cells were washed with PBS, normalized to OD₆₀₀ = 1 to ensure equal bacterial numbers, and TTC reduction was quantified by measuring optical density at 496 nm. Treatment with 0.5 mM indole resulted in approximately 10-fold reduction in TTC reduction relative to DMSO-treated bacteria, indicating marked impairment of respiratory activity **(SFig. 3B)**. These data support the transcriptomic findings and demonstrate that indole inhibits *C. jejuni* growth, at least in part, by suppressing cellular respiration.

In addition to its effects on respiratory gene transcript abundance, indole resulted lower abundance of transcripts from genes of the TCA cycle genes and the *ackA*-*pta* acetate switch pathway, which are important for ATP generation and metabolic flux in *C. jejuni*. To determine whether indole affects cellular energy status, we measured intracellular ATP levels. Wild type bacteria were grown for nine hours in different concentrations of indole, normalized to OD₆₀₀ = 1, and intracellular ATP was quantified using a commercial luminescence-based assay. Consistent with impaired respiration, 0.5 mM indole caused an approximately three-fold reduction in intracellular ATP levels compared with controls **(SFig. 3C)**, consistent with previous findings showing that indole reduces intracellular ATP levels in *Pseudomonas putida* (38). This suggests a metabolic mechanism underlying indole-mediated growth inhibition.

In our transcriptomic analysis, abundance of mRNA from several oxidative stress regulatory genes was significantly elevated in the presence of indole, including *perR* (peroxide-sensing regulator), *katA* (catalase), *aphC* (alkyl hydroperoxide reductase), and *sodB* (Superoxide dismutase B), showing approximately two- to nine-fold increase in abundance. In addition, transcripts from iron-regulated genes, cj0175c (*cfbpA*), cj0174c (*cfbpB*), cj1659 (*p19*), and the associated ABC transporter genes cj1660–cj1663, and the inorganic phosphate transporter operon *pstSCAB*, were elevated by roughly six- to eight-fold following indole exposure. These genes are induced under acidic conditions in *C. jejuni*. Furthermore, transcripts from cj0414 and cj0415, which encode gluconate dehydrogenase subunits, exhibited a six- to sevenfold increase in expression in the presence of indole. These genes promote *C. jejuni* survival under acidic conditions **(sTable 2).**

Consistent with these observations, our transcriptomic data reveal greatly increased abundance of mRNA from acid-adaptation and stress-response genes in *C. jejuni* following indole exposure, suggesting induction of a response associated with intracellular acidification. We therefore hypothesize that indole inhibits *C. jejuni* growth by reducing cytoplasmic pH. To test this, wild-type *C. jejuni* was grown for nine hours in the presence or absence of indole, washed with PBS, and normalized to OD₆₀₀ = 1. Cells were stained with the pH-sensitive fluorescent dye CMFDA, and fluorescence emission was measured at 520 nm with dual excitation at 450 nm and 490 nm. Intracellular pH was calculated using a standard calibration curve generated under defined pH conditions. Indole treatment resulted in a significant reduction in intracellular cytoplasmic pH compared with untreated controls. While control cells-maintained a near-neutral cytoplasmic pH approximately 7.2, indole-exposed bacteria exhibited a decreased intracellular pH of approximately 6 **(SFig. 3D)**. Thus, indole disrupts intracellular pH homeostasis in *C. jejuni*, providing an additional mechanistic explanation for indole-mediated growth inhibition, likely through metabolic and respiratory stress associated with cytoplasmic acidification.

To test the requirement for lactate-dependent growth of *C. jejuni* in the DSS-treated mouse, we performed a competitive colonization assay in DSS-treated mice using a 1:1 mixture (10⁸ CFU total) of wild-type and *lctP::kan* mutant strain. Colonization was quantified in colonic tissue at day three post-infection. The *lctP::kan* mutant exhibited greater than 100-fold fitness defect relative to the wild-type strain, demonstrating a marked competitive disadvantage in vivo **(Fig. 4F)**. These findings indicate that lactate uptake via LctP is critical for *C. jejuni* fitness during DSS-induced colitis. Together, our data support a model in which inflammation-associated oxygenation and lactate accumulation create a metabolic niche that favors *C. jejuni* growth through oxygen-enhanced lactate metabolism, while indole acts to suppress this pathway.

Nitrate respiration, fueled by host-derived nitrate produced through iNOS-dependent nitric oxide synthesis, provides a growth advantage to several enteric pathogens in the inflamed gut. Unlike *E. coli* and *S. enterica* serovar Typhimurium, which encode multiple nitrate reductases(12, 39), *C. jejuni* possesses a single periplasmic nitrate reductase encoded by the *napAGHBLD* operon(39), and indole negatively impacts transcript abundance from this operon. Nitrate respiration in *C. jejuni* is required for efficient commensal colonization in the chicken intestine, while the contribution of nitrate respiration to *C. jejuni* fitness during inflammatory infection remains poorly defined(40). We generated a nitrate reductase–deficient mutant (*napA::cm*), which is impaired in nitrate respiration, and performed a competitive colonization assay in DSS-treated mice using a 1:1 mixture (10⁸ CFU total) of wild-type and *napA::cm* bacteria. Bacterial burdens were quantified in colonic tissue at day three post-infection. The *napA::cm* mutant exhibited an approximately 100-fold fitness defect relative to the wild-type strain, indicating a marked competitive disadvantage in vivo **(Fig. 4F)**. We observed a ∼five-fold increase in colonic *iNOS* transcription following DSS treatment and an approximately 10-fold increase during infection **(SFig. 3E)**, with elevated expression of iNOS detected with immunohistochemistry; this is observed mainly in colonic epithelial cells in DSS uninfected and infected tissue **(SFig. 3F)**. These findings suggest that inflammation enhances the availability of alternative electron acceptors in the inflamed gut and that *C. jejuni* relies on nitrate respiration to maximize fitness in this environment.

We also investigated the *ackA/pta* pathway of ATP generation and its role in *C. jejuni* growth in the inflamed gut., as its requirement for commensal, non-inflammatory colonization of poultry has already been demonstrated(37). After inoculating DSS-treated mice with 1:1 mixture (10⁸ CFU total) of wild-type and *ackA/pta*::cm bacteria, the mutant demonstrated approximately 100-fold competitive defect after three days **(Fig. 4F)**.

Given our *in vitro* finding that indole inhibits *C. jejuni* growth, and that several genes whose expression is negatively affected by indole are required for fitness in the inflamed gut, we next evaluated whether indole administration could limit bacterial colonization *in vivo*. DSS-treated mice were orally infected with *C. jejuni* (10⁸ CFU) as described above and, following infection, one group received oral indole (150 mg/kg of body weight, prepared in DMSO) once each of the two days post-infection, while the control group received DMSO alone **(Fig. 5A)**. Mice treated with indole exhibited an approximately 1,000-fold reduction in *C. jejuni* colonization compared with control mice treated with DMSO alone **(Fig. 5B)**. Indole treatment did not significantly alter histological scores or inflammatory parameters compared with DMSO-treated infected mice. **(Fig. 5C)**, indicating that the indole has a direct inhibitory effect on bacterial growth and not on intestinal inflammation.

**Figure 5.**
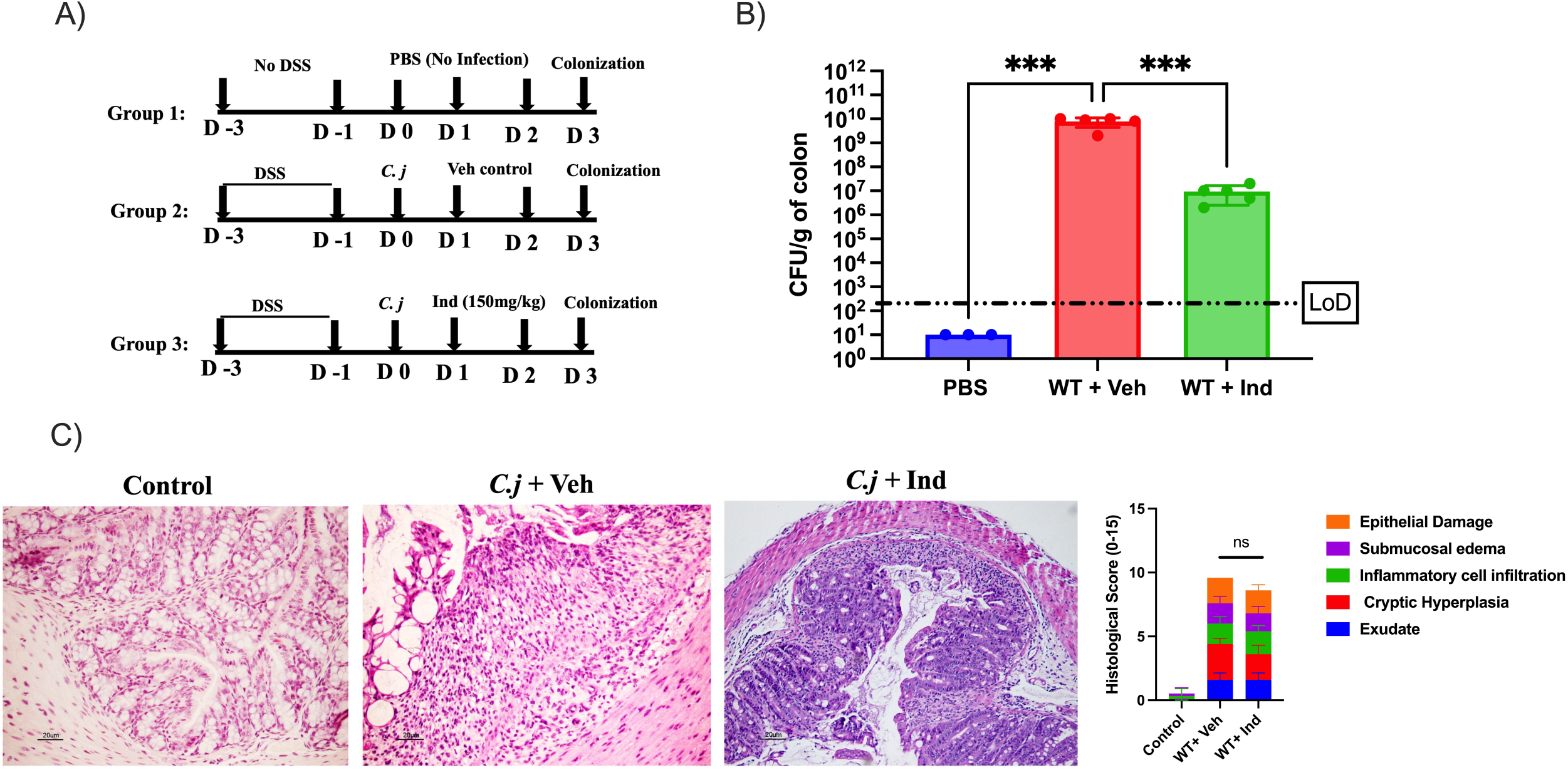
Indole treatment reduces *C. jejuni* colonization in vivo. (A) Schematic diagram of the experimental design for indole or vehicle (control) treatment following *C. jejuni* infection. (B) Comparative colonization of *C. jejuni* in colon tissues between indole-treated and vehicle-treated groups on day 3 post-infection, determined by colony-forming unit (CFU) counts (n = 5). (C) Colonic inflammatory responses were evaluated by histological analysis and histological scoring of hematoxylin and eosin (H&E)-stained sections on day 3 post-infection. Representative images (H&E) and histological scores are shown. Data are presented as mean ± SE. Statistical analysis was performed using one-way ANOVA. *P* < 0.05; **P** < 0.01; ***P*** < 0.001; **P** < 0.0001.

### Probiotic indole production inhibits *C. jejuni* growth *in vivo*

Among other bacteria, the probiotic *Escherichia coli* strain Nissle 1917 (EcN) produces indole from L-tryptophan via tryptophanase A (*tnaA*)(41). To examine the impact of bacterial indole production on *C. jejuni* fitness, we grew co-cultures of *C. jejuni* 11168 and either wild type or *tnaA* mutant EcN in Mueller–Hinton broth (MHB) supplemented with L-tryptophan (2 mM). After 24 hours, *C. jejuni* CFUs were quantified, and indole concentrations in culture supernatants were measured using the Kovac assay. Co-culture with wild-type EcN in the presence of tryptophan resulted in significant suppression of *C. jejuni* growth, coinciding with indole accumulation (∼1 mM) in the supernatant **(Fig. 6A)**. In contrast, co-culture with the *tnaA::cm* mutant did not impair *C. jejuni* growth, and indole was undetectable in the supernatant. Growth of *C. jejuni* alone in tryptophan-containing medium was unaffected, confirming that growth inhibition was dependent on EcN-derived indole production rather than tryptophan supplementation itself **(Fig. 6B)**. These findings demonstrate that microbially-derived indole production is sufficient to suppress *C. jejuni* growth *in vitro*.

**Figure 6.**
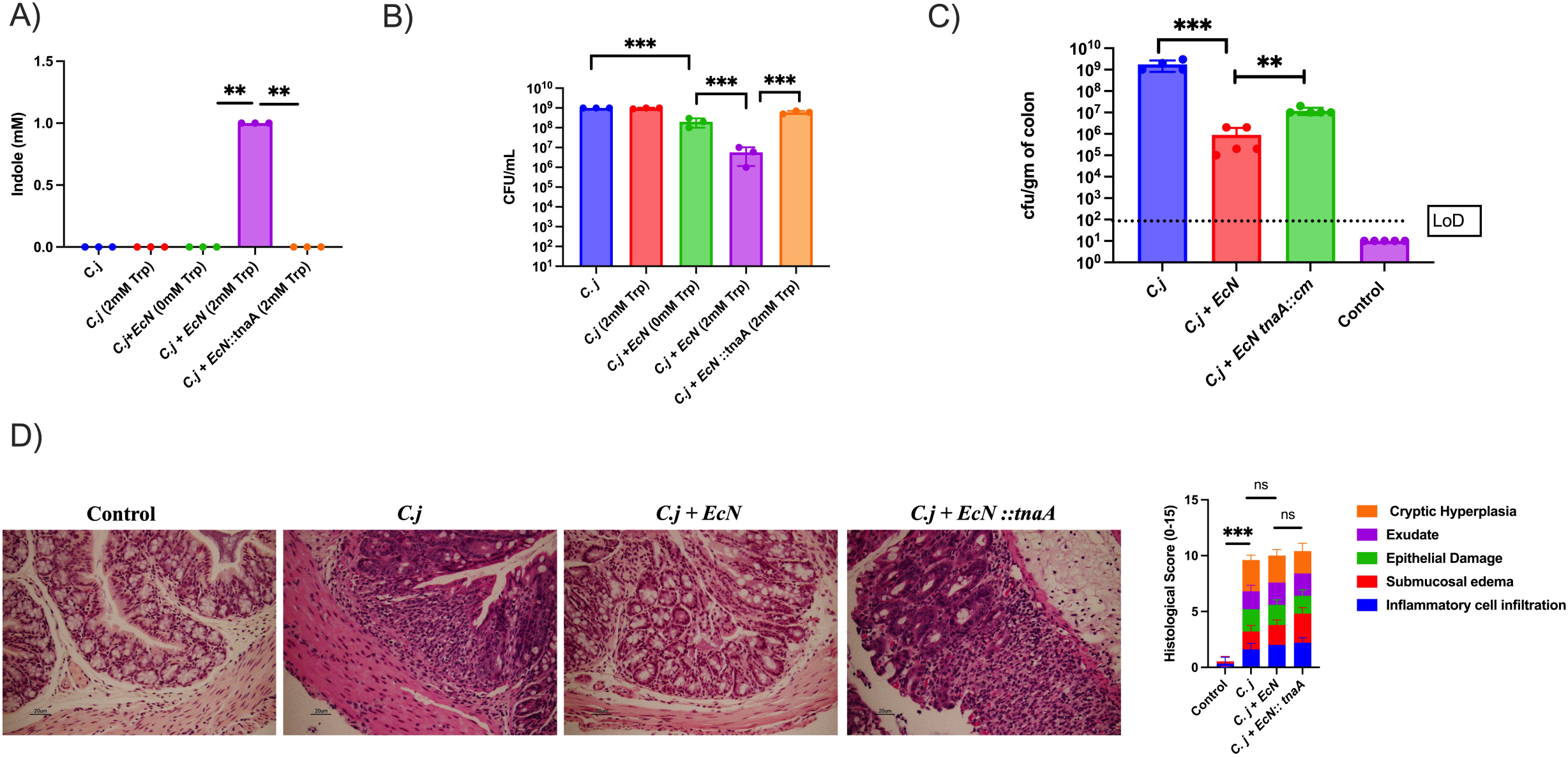
Indole produced by probiotic *E. coli* Nissle 1917 (EcN) inhibits *C. jejuni* colonization *invitro* and *invivo*. (A) Indole production and (B) *C. jejuni* growth were measured after 24 h in Mueller–Hinton broth (MHB) supplemented with and without tryptophan (2 mM). Cultures included *C. jejuni* alone, *C. jejuni* co-cultured with WT EcN, and *C. jejuni* co-cultured with the EcN *tnaA::cm* mutant. Indole levels were quantified, and *C. jejuni* growth was determined by colony-forming unit (CFU) counts. (C) Comparative colonization of *C. jejuni* in colon tissues was determined by CFU counts among mice infected with *C. jejuni* alone, co-infected with *C. jejuni* and WT EcN, or co-infected with *C. jejuni* and the EcN *tnaA::cm* mutant. (D) Colonic inflammatory responses were evaluated by histological analysis and histological scoring of hematoxylin and eosin (H&E)-stained sections on day 3 post-infection among three groups: control, *C. jejuni* + EcN, and *C. jejuni* + EcN *tnaA::cm*. Representative images (H&E) and histological scores are shown. *Error bars represent SD. Statistical analysis was done by unpaired two-tailed t test. *P < 0.05; **P < 0.01; ***P < 0.001; ****P < 0.000*.

We also investigated whether indole-producing EcN could suppress *C. jejuni* colonization during infection. DSS-treated mice were orally infected with *C. jejuni* alone or co-infected at a 1:1 ratio (10⁸ CFU total) with either wild-type EcN or the *EcN tnaA::cm* mutant incapable of producing indole. Colonic colonization was assessed at day three post-infection. Co-infection with wild-type *EcN* resulted in an approximately 100-fold reduction in *C. jejuni* colonization compared with co-infection with the *tnaA::cm* mutant. In contrast, *C. jejuni* grew to high levels when co-infected with the EcN *tnaA::cm* strain which cannot produce indole. Infection with *C. jejuni* alone yielded colonization levels comparable to those observed in the *tnaA::cm* co-infection group **(Fig. 6C)**. Histological scores and other inflammatory parameters were not altered across the three infected groups suggesting that the impact of indole production was on the *C. jejuni* growth *per se* and not on the host **(Fig. 6D).**

Our findings support the hypothesis that metabolic competition mediated by probiotic derived indole is a mechanism of colonization resistance during intestinal inflammation.

## Discussion

The gastrointestinal (GI) tract is densely populated by microbial communities that maintain colonization resistance and protect the host from enteric pathogens(42). Resident microbiota restrict pathogen growth by limiting nutrient availability and maintaining a hypoxic intestinal environment, partly through microbiota-derived short-chain fatty acids such as butyrate, metabolism of which preserves epithelial hypoxia(10). Pathogenic bacteria disrupt this barrier by inducing intestinal inflammation, creating conditions favorable for their growth. Inflammation remodels the intestinal ecosystem by altering epithelial metabolism, oxygen availability, nutrient flux, and microbial composition, a mechanism well described for facultative anaerobic Enterobacteriaceae such as *S.* Typhimurium and *E. coli*, which exploit inflammation-associated electron acceptors and increased oxygen to outcompete obligate anaerobic commensals(10).

In previous work with a ferret infection model, we showed that intestinal inflammation supports growth of *C. jejuni* by increasing epithelial oxygenation and shifting metabolism towards one that elevates lactate levels, which is used as a growth substrate by the pathogen(3). In the current study we generated a similar inflammatory environment in the mouse using DSS-induced colitis and demonstrated that transient inflammation alone is sufficient to overcome colonization resistance and allow robust *C. jejuni* growth in conventional mice without antibiotic treatment or host genetic modification. This outcome is consistent with the widely used IL10–deficient mouse model, in which constitutive intestinal inflammation results in high susceptibility to *C. jejuni* colonization; these two models provide further evidence that intestinal inflammation is a key determinant of *C. jejuni* fitness and disease progression(4).

Inflammation not only facilitates bacterial growth but is also intensified by infection, establishing a feed-forward interaction between host inflammatory responses and pathogen fitness. These findings may help explain clinical observations that *Campylobacter* species are frequently isolated from patients with active colitis, and acute enteric infection is known to trigger disease flares in individuals with inflammatory bowel disease (IBD) (43). Additionally, epidemiological studies have reported that acute *Campylobacter* infection is associated with an increased long-term risk of developing IBD (44).

This mouse model provides a simple, reproducible, and cost-effective platform to study *C. jejuni* pathogenicity. It is well suited for identifying bacterial physiological traits and fitness determinants specifically required under inflammatory conditions, and to address host traits that contribute to pathogenicity in the gut.

In patients with inflammatory bowel disease, particularly ulcerative colitis, epithelial metabolic changes like those observed in DSS-induced colitis have been reported (7). DSS treatment induces ER stress–mediated epithelial injury, leading to enhanced repair responses characterized by crypt elongation, proliferation of transit-amplifying cells, and reduced numbers of terminally differentiated cells such as goblet cells, resulting in thinning of the mucus layer(45). These histological changes were also observed in our model and were exacerbated following *C jejuni* infection. Terminally differentiated colonocytes typically rely on β-oxidation of fatty acids, consuming high levels of oxygen, whereas proliferating epithelial cells preferentially use glycolysis, producing lactate with lower oxygen consumption (10). Consistent with previous reports showing elevated luminal lactate during colitis, this metabolic shift reduces epithelial oxygen use and increases oxygen and lactate availability at the mucosal surface (8, 9). In our model, epithelial oxygenation increased after DSS treatment and further after *C. jejuni* infection, suggesting higher oxygen levels in the gut lumen. Lactate is also critical for *C. jejuni* growth under inflammatory conditions. Because *C. jejuni* is a microaerophile requiring low oxygen tension (2–10%) for respiration, inflammation-associated epithelial oxygenation likely creates a microaerobic niche while simultaneously increasing lactate availability, providing both an electron acceptor and a carbon source that support *C. jejuni* growth in the inflamed intestine. Previous studies have shown similar growth physiology in *S. enterica* Typhimurium infection (11).

Host-derived nitrate serves as an alternative electron acceptor for facultative anaerobic bacteria such as *E. coli* and *S*. *enterica* Typhimurium during intestinal inflammation (10, 12, 39). During colitis, the host inflammatory response generates reactive nitrogen and oxygen species, and inducible nitric oxide synthase (iNOS) is elevated, leading to increased nitric oxide production in the gut. Nitric oxide reacts with superoxide to form peroxynitrite, which can subsequently generate nitrate, thereby increasing luminal nitrate availability (10, 12). We detected elevated iNOS expression in both DSS-treated and *C. jejuni*–infected colonic tissue, suggesting increased nitrate levels in the inflamed intestine. Additionally, the nitrate reductase mutant showed a competitive colonization defect, indicating that nitrate respiration contributes to *C. jejuni* fitness in the inflamed gut. Overall, our study suggests that luminal lactate availability, epithelial oxygenation, and nitrate levels are key factors supporting bacterial growth in the inflamed gut. Diet can strongly influence these parameters; for example, long-term high-fat diets increase epithelial oxygenation and nitrate availability, thereby promoting growth of Enterobacteriaceae such as *Escherichia coli* and *Salmonella enterica* serovar Typhimurium (46). However, little is known about how dietary factors affect the pathogenesis of *C. jejuni*. Our findings suggest that diet-driven changes in gut metabolism and redox conditions may influence *C. jejuni* colonization and disease progression, highlighting an important area for future investigation.

We also identified luminal indole as important in this model. DSS treatment and *C. jejuni* infection significantly reduced luminal indole levels, correlating with decreased abundance of indole- producing members of the microbiota, consistent with reports showing reduced indole concentrations during intestinal inflammation and in patients with inflammatory bowel disease (47). Indole inhibited *C. jejuni* growth both *in vitro* and *in vivo*, with transcriptomic analysis indicating that indole results in reduced abundance of transcripts for aerobic respiration, lactate utilization, nitrate respiration and acetate switch pathways, all critical determinants of bacterial fitness in the inflamed intestine.

Although we generated inflammation experimentally leading to an altered, less-indole-producing microbiota, presumably *C. jejuni* must overcome indole-mediated colonization resistance during natural infection to establish disease. Microbiota from humans infected with *Campylobacter* have similar changes to what we observed with DSS-treatment in the mouse such as lower levels of *Firmicutes*, which include indole-producing *Clostridia* species (48). To draw solid conclusions about if and how *C. jejuni* alters the gut microbiota in a way that reduces indole-producing bacteria or otherwise relieves metabolic inhibition requires further study. In general, understanding how *C. jejuni* modulates the intestinal metabolic environment to overcome indole-dependent resistance is an important area for future investigation.

With increasing emergence of multidrug-resistant *C. jejuni* strains, alternative therapeutic strategies are needed (2). Our results suggest that a metabolite-based approach to limit infection could be successful. Consistent with this idea, recent studies have reported that indole and synthetic indole-derived compounds exhibit antimicrobial or anti-virulence activity against a broad range of Gram-positive and Gram-negative pathogens, supporting this approach for therapeutic development against multidrug-resistant bacteria (49, 50).

Collectively, this study reframes *C. jejuni* pathogenesis within the ecology of the inflamed gut. Less dependent on virulence determinants such as type III secretion effectors or exotoxins that commandeer host functions, *C. jejuni* exploits inflammation-induced oxygenation, nitrate availability, and glycolytic products to fuel respiratory metabolism. Concurrent depletion of indole-producing commensals removes a critical metabolic constraint on this growth. By integrating host metabolic remodeling, microbial community restructuring, and pathogen bioenergetics, our study provides a unified model explaining how inflammation converts the colon into a permissive niche for *C. jejuni*. These findings also highlight microbial metabolite restoration—particularly indole—as a rational therapeutic strategy to limit pathogen growth without broadly suppressing inflammation or disrupting the microbiota.

## Materials and Methods

### Bacterial strains and media

All strains and plasmids used in the study are listed in **Table S1**. All *C. jejuni* strains were routinely grown under microaerobic conditions (85% N2,10% CO2, 5% O2) on Mueller-Hinton (MH) agar or in broth under necessary antibiotics at the following concentrations: kanamycin, 50 μg ml−1; trimethoprim (TMP), 5 μg ml−1. *C. jejuni* strains were stored in MH broth with 20% glycerol at −80°C. *Escherichia coli* Nissle 1917 (EcN) were routinely grown under aerobic condition at 37^0^C in Luria Broth agar (LB).

Construction of C. jejuni napA::cm, ackA::pta::cm and E. coli Nissle 1917 tnaA::cm insertion mutant.

The *C. jejuni napA::cm* and *ackA::pta::cm* mutants were constructed by inserting a chloramphenicol resistance cassette into the gene as previously described. Briefly, 500 bp of both upstream and downstream regions of the *napA* and *ackA* locus was amplified by PCR with primers containing *Sma*I restriction sites. The amplified upstream and downstream regions were joined by PCR. The 1,000-bp amplified PCR product was cloned into the plasmid pGEM-T Easy. The *Campylobacter* chloramphenicol resistance cassette was excised from pRY109 using *Sma*I and ligated between the upstream and downstream cloned regions of the pGEM-T construct. The plasmid was electroporated into the wild type strain and grown on MH agar overnight at 37°C under microaerobic conditions. Cells were then plated and grown on MH agar containing chloramphenicol for 2 to 3 days, and successful integration of ***napA::cm*** and ***ackA::pta::cm*** into the chromosome was confirmed by PCR (3).

The *E. coli* Nissle 1917 ***tnaA::cm*** mutant was constructed by inserting a chloramphenicol resistance cassette into the open reading frame of ***tnaA*** using λ Red recombineering. Deletion–insertion alleles were generated by PCR amplification of the chloramphenicol resistance cassette from pKD3 (42) using primers containing 50-bp homology arms targeting the gene of interest. The first and last three codons of the open reading frame were retained. The resulting PCR products were introduced into EcN harboring pKD46 (43) by electroporation. Following electroporation, mutants were selected on LB agar containing chloramphenicol (15 μg/ml), and the mutation was confirmed by PCR(51).

### Growth analysis in MH broth

*C. jejuni* wild-type strain was recovered from freezer stocks and cultured on Mueller–Hinton (MH) agar supplemented with appropriate antibiotics for 24 h under microaerobic conditions, followed by restreaking and incubation for an additional 16 h. Bacteria were then resuspended in MH broth and inoculated into 25 mL of Mueller–Hinton broth containing indole (0.25, 0.5, or 1 mM) or vehicle control (DMSO) at an initial OD₆₀₀ of ∼0.05. Cultures were incubated at 37 °C with shaking (100 rpm) under microaerobic conditions. Bacterial growth was monitored over 30 h by enumerating colony-forming units (CFUs) at indicated time points.

For coculture assays, *Escherichia coli* Nissle 1917 (EcN) and *Campylobacter jejuni* wild-type (WT) strains were inoculated individually or together into 25 mL Mueller–Hinton broth (MHB) supplemented with L-tryptophan (2 mM) or vehicle control (DMSO), with an initial OD₆₀₀ of ∼0.05 for each strain. Cultures were incubated at 37 °C with shaking (100 rpm) under microaerobic conditions for 24 h, after which *C. jejuni* growth was quantified by enumerating colony-forming units (CFUs) on selective Campylobacter agar plates (cefoperazone (40 µg/ml), cycloheximide (100 µg/ml), trimethoprim (5 µg/ml), and vancomycin (100 µg/ml). Indole concentrations in culture supernatants were measured using a modified Kovács assay (52). Briefly, cultures were centrifuged at 15,000 rpm for 15 min, and 100 μL of supernatant was incubated in triplicate with 150 μL Kovács reagent (Sigma-Aldrich) for up to 30 min at room temperature. The resulting chromophore was quantified spectrophotometrically at 530 nm, and indole concentrations were determined using standard curves prepared in MHB.

### Mouse infections

All animal experiments were approved by the Michigan State University Institutional Animal Care and Use Committee (IACUC). Approval No. PROTO202400286).

1. Specific pathogen-free male and female C57BL/6J mice (7–8 weeks old; The Jackson Laboratory) were randomly assigned to four groups (n = 5 per group): (i) no DSS, no *C. jejuni* infection; (ii) no DSS, *C. jejuni* infection; (iii) DSS treatment only; and (iv) DSS treatment with *C. jejuni* infection. For DSS treatment, mice received 3% dextran sodium sulfate (DSS, MP Biomedicals Colitis grade (MW36,000 - 50,000 Da) in drinking water for 3 days. On day 4, mice were deprived of food and water for 2–3 h prior to infection and then orally gavaged with *C. jejuni* wild-type strain (10^8^ CFU) in 5% sodium bicarbonate to neutralize gastric acid; DSS was subsequently replaced with regular drinking water. Control mice received PBS with sodium bicarbonate. At 1 and 3 days postinfection, mice were euthanized, and fecal samples and tissues (cecum, small intestine, and proximal and distal colon) were collected to assess bacterial burden. Samples were homogenized in sterile phosphate-buffered saline (PBS), serially diluted, and plated on selective Campylobacter agar plates. Plates were incubated for 48 h under microaerophilic conditions, and colonies were enumerated(3).
2. For *invivo* competition assays, DSS-treated female mice were orally infected with a 1:1 mixture (total 10^8^ CFU) of *C. jejuni* wild-type and mutant strains (*lctP::kan*, *napA::cm*, or *ackA::pta::cm*). At 3 days postinfection, colonic tissues were collected, homogenized, and serially diluted in sterile PBS. Samples were plated on Campylobacter-selective agar with or without selective antibiotics (kanamycin, 50 μg/mL; chloramphenicol, 15 μg/mL) to differentiate strains. Competitive indices were calculated as previously described.
3. For indole treatment on *C. jejuni* colonization *invivo*, female mice were assigned to three groups: (i) no DSS, no infection; (ii) DSS treatment with *C. jejuni* infection plus indole; and (iii) DSS treatment with *C. jejuni* infection plus vehicle control. Indole (Sigma-Aldrich) was administered by oral gavage twice daily on days 2 and 3 postinfection at a total dose of 150 mg·kg^−1^·day^−1^ in a vehicle consisting of DMSO/PEG400/5% citric acid (1:4.5:4.5)(28). At 3 days postinfection, colonic tissues were collected, homogenized in sterile PBS, and serially diluted. Bacterial burden was determined by plating on Campylobacter-selective Mueller–Hinton agar.
4. For *invivo* coinfection assays, DSS-treated female mice were orally inoculated with 1 × 10^8 CFU of a 1:1 mixture of *C. jejuni* and *Escherichia coli* Nissle 1917 or its *tnaA::cm* mutant. At 3 days postinfection, colonic tissues were collected, homogenized in sterile PBS, and serially diluted. *C. jejuni* colonization levels were determined by plating on Campylobacter-selective Mueller–Hinton agar.

### Histology and Immunohistochemistry

Colonic tissues were fixed in 10% neutral-buffered formalin and transferred to 60% ethanol prior to processing. Samples were sectioned at 5 μm, mounted on glass slides, and stained with hematoxylin and eosin (H&E) and Alcian blue to assess goblet cells. Histopathological evaluation was performed in a blinded manner using a semiquantitative scoring system based on epithelial damage, inflammatory cell infiltration, goblet cell depletion, crypt hyperplasia, and crypt abscess formation (0, none; 1, minimal; 2, mild; 3, moderate; 4, marked; 5, severe). Localization of *Campylobacter jejuni* in colonic tissues was assessed by immunohistochemistry using a polyclonal anti-*C. jejuni* primary antibody. Cell proliferation and inflammatory responses were evaluated by staining for Ki67^+^ and NOS2 using specific anti-mouse primary antibodies. All histological and immunohistochemical analyses were performed at the Investigative HistoPathology Laboratory, Department of Physiology, Michigan State University.

### Hypoxia staining

Tissue hypoxia was assessed using pimonidazole labeling(8). Mice were administered pimonidazole HCl (60 mg/kg, i.p.; Hypoxyprobe-1 kit, Hypoxyprobe) 1 h prior to euthanasia. Colonic tissues were fixed in 10% phosphate-buffered formalin, paraffin embedded and processed for immunofluorescence staining. Sections were blocked using a mouse-on-mouse blocking reagent (Vector Laboratories) and incubated with a mouse anti-pimonidazole monoclonal antibody (MAb 4.3.11.3). Nuclei were counterstained with DAPI using SlowFade Gold mounting medium. Hypoxia was scored in a blinded manner based on epithelial staining intensity and distribution (0, none; 1, mild focal; 2, moderate multifocal; 3, intense diffuse) as previously described(8). Representative images were acquired using a microscope, and image brightness was adjusted uniformly across samples.

### RNA-seq and Real-time PCR

*C. jejuni* strain 11168 was cultured in Mueller–Hinton broth supplemented with 0.5 mM indole or vehicle control (DMSO) for 9 h. Total RNA was extracted using TRIzol reagent followed by purification with the RNeasy Mini Kit (Qiagen)(53, 54). Residual genomic DNA was removed by TURBO DNase treatment (Invitrogen). Libraries were sequenced at SeqCenter, Pittsburgh, USA, generating 12 million paired-end reads (2 × 150 bp) per sample (n = 3). The data were analyzed using Proseq v2.0 software. Detailed analysis methods are available at the project repository: https://github.com/snandiDS/prokseq-v2.0.

For RT-PCR analysis, first-strand cDNA was synthesized using SuperScript III First-Strand Synthesis SuperMix (Invitrogen). Quantitative real-time PCR was performed using SYBR Green chemistry, and gene expression levels were calculated using the 2^^−ΔΔCt^ method as previously described(54). Primer sequences for all genes analyzed are provided in Table S2.

Proinflammatory and anti-inflammatory cytokine expression in colonic tissues from DSS-treated, infected, and control mice was assessed as previously described. Flash-frozen tissues were homogenized in TRIzol reagent, and total RNA was extracted followed by purification using the RNeasy Mini Kit (Qiagen). Residual genomic DNA was removed by TURBO DNase treatment (Invitrogen). First-strand cDNA was synthesized, and quantitative real-time PCR was performed using SYBR Green chemistry to measure expression of *Tnf -α*, *Il6*, *Cxcl1* (KC), *Nos2*, and *Il10*. Relative gene expression was calculated using the 2^−ΔΔCt method as described above. Primer sequences are provided in Table S2.

### 16S rRNA sequencing

Colonic contents were collected and microbial DNA was extracted using the DNeasy PowerSoil Pro kit (Qiagen) according to the manufacturer’s instructions(3). Amplicon libraries targeting the V3–V4 regions of the 16S rRNA gene were generated using the Quick-16S Library Prep Kit (Zymo Research) with forward primers 341F (CCTACGGGDGGCWGCAG) and its variant (CCTAYGGGGYGCWGCAG), and reverse primer 806R (GACTACNVGGGTMTCTAATCC). Libraries were purified, normalized, and sequenced on an Illumina NextSeq 2000 platform (2 × 301 bp). Base calling, demultiplexing, and adapter trimming were performed using Illumina BCL Convert (v4.2.4), followed by quality filtering (minimum read length >150 bp; poly-N <10; poly-G <150). Sequence processing was conducted in QIIME 2 (v2024.5) using the DADA2 plugin for denoising, chimera removal, and amplicon sequence variant (ASV) inference, with reads truncated based on quality scores (Phred ≥24), yielding final lengths of ∼275–290 bp. Taxonomic classification was performed using a Naïve Bayes classifier trained on the Greengenes2 database (release 2024.9.2) trimmed to the V3–V4 region using experimental primers. Alpha diversity (Observed ASVs, Shannon index, Chao1) and beta diversity (Bray–Curtis and Jaccard distances) were calculated using the QIIME 2 core-metrics-phylogenetic pipeline, with group differences assessed by Kruskal–Wallis test and visualized by principal coordinates analysis (PCoA).

## Metabolite Analysis

Indoles and tryptophan-derived metabolites were quantified from colonic contents collected from infected and uninfected groups at day 3 postinfection using liquid chromatography–tandem mass spectrometry (LC-MS/MS). Samples were extracted in 1 mL of 80:20 (vol/vol) methanol/water containing internal standards, followed by centrifugation to remove debris. Supernatants were subjected to LC-MS/MS analysis.

A 2-μL aliquot of each sample was injected onto an Acquity HSS-T3 UPLC column (2.1 × 150 mm; Waters) maintained at 50 °C and separated using an Acquity UPLC system (Waters) coupled to a Xevo TQ-XS triple quadrupole mass spectrometer (Waters). Chromatographic separation was performed at a flow rate of 0.3 mL/min using a 14-min gradient with mobile phase A (water containing 0.6% formic acid) and mobile phase B (methanol containing 0.6% formic acid). Initial conditions (98% A/2% B) were held for 1 min, followed by a linear gradient to 95% B over 10 min, maintained at 95% B until 11 min, and then returned to initial conditions (98% A) at 11.01 min and equilibrated until 14 min.

Analytes were detected by electrospray ionization in both positive and negative ion modes using multiple reaction monitoring. Instrument parameters were optimized for each metabolite. Data acquisition and quantification were performed using TargetLynx software (Waters MassLynx platform).

### ATP measurement

Intracellular ATP levels were measured using the BacTiter-Glo assay kit (Promega) according to the manufacturer’s instructions. *C. jejuni* wild-type strain was cultured for 9 h in 25 mL Mueller–Hinton broth supplemented with 0.5 mM indole or vehicle control (DMSO), starting at an initial OD₆₀₀ of 0.05. Cultures were harvested by centrifugation, washed twice with PBS, and normalized to an OD₆₀₀ of 1.0. ATP levels were then quantified using the BacTiter-Glo reagent, and concentrations were determined based on a standard curve(54).

### Respiration Assays

*Campylobacter jejuni* wild-type strain was cultured for 9 h in 25 mL Mueller–Hinton broth supplemented with 0.5 mM indole or vehicle control (DMSO), starting at an initial OD₆₀₀ of 0.05. Cultures were harvested by centrifugation, washed twice with PBS, and normalized to an OD₆₀₀ of 1.0. For respiration measurements, 135 μL of the cell suspension was mixed with 15 μL of 1% (wt/vol) 2,3,5-triphenyltetrazolium chloride (TTC). Control reactions lacking substrate (water only) were included as blanks. Samples were incubated for 1 h under microaerophilic conditions at 42 °C, and TTC reduction was quantified by measuring absorbance at 496 nm.(54).

### Statistical Analysis

Statistical differences across metadata categories, such as infection status, were evaluated using permutational multivariate analysis of variance (PERMANOVA) with 999 permutations, including pairwise post-hoc testing. Genus-level differential abundance testing was performed using Analysis of Compositions of Microbiomes with Bias Correction (ANCOM-BC^3^). Holm’s method corrected *p*-values for multiple comparisons, and significant taxa (adjusted *p* < 0.05) were used for further analysis. All microbiota analysis graphs and plots were generated using R (version 4.5.1^4^) with the packages ggplot2(56) and phyloseq(57).

## Acknowledgments

This work was supported by the Michigan State University Rudolph Hugh Endowment (to V.J.D.). We would like to thank Prof. Christine M. Szymanski, Department of Microbiology, University of Georgia, Athens, Georgia, USA and Prof. Shannon Manning, Department of Microbiology, Genetics and Immunology, Michigan State University for providing *C. jejuni* strains.

## Supplementary

**Fig. S1. *C. jejuni* colonization in DSS-treated mice.** (A) Colonization of *C. jejuni* in different intestinal segments of DSS-treated mice, quantified by CFU counts at day 3 postinfection. (B) Colonization in proximal and distal colon regions at day 3 postinfection. (C) Sex-based comparison of *C. jejuni* colonization in colon samples from female and male mice at day 3 postinfection. Data represent mean ± SD (n = 5 per group). Statistical significance was determined using an unpaired two-tailed t test. *P < 0.05, **P < 0.01, ***P < 0.001, ****P < 0.0001.

**Fig. S2. Gut microbiota analysis in DSS-treated mice.** (A) Alpha diversity metrics (Chao1 richness, observed amplicon sequence variants [ASVs], and Shannon index) shown as box plots with data rarefied to the minimum sequencing depth. Statistical comparisons were performed using the Kruskal–Wallis test. (B) Principal coordinates analysis (PCoA) based on Jaccard distances showing separation between uninfected controls and DSS-treated groups. Significance was assessed by PERMANOVA (999 permutations) with pairwise post hoc comparisons (*P < 0.05, **P < 0.01, ***P < 0.001). (C) Beta diversity analysis showing Jaccard distances between groups, highlighting differences between uninfected controls and DSS-treated mice. (D) Differential abundance of bacterial genera in DSS-treated groups (untreated and infected) relative to uninfected controls. Log fold change (LFC) analysis identifies significantly enriched (red) and depleted (purple) taxa.

**Fig. S3. Effects of indole on *C. jejuni* physiology and host responses.** (A) Growth of *C. jejuni* isolates in the presence of varying concentrations of indole, measured by CFU counts after 24 h. (B) Respiratory activity in the presence of indole (0.5 mM) or vehicle control (DMSO), assessed by TTC assay after 9 h (n = 3). (C) Intracellular ATP levels measured after 9 h in the presence or absence of indole (0.5 mM) (n = 3). (D) Cytoplasmic pH measured after 9 h in the presence or absence of indole (n = 3). (E) Relative *Nos2* gene expression in colon tissues from DSS-treated uninfected and *C. jejuni*-infected mice compared with PBS controls, determined by qRT-PCR (n = 5). (F) iNOS protein expression in colon tissues at day 3 postinfection, assessed by immunohistochemistry using anti-NOS2 antibody. Representative images (H&E) and histological scores are shown.

## Author contributions

R.S. and V.J.D. designed research; R.S., E.O.,R.M.L,V.J.D performed research; R.S.,B.B.,C.K.Z.,V.J.D analyzed data, R.S., B.B, P.S.,V.J.D. wrote the manuscript, R.S, P.S.; V.J.D edited the manuscript.

