## Supplementary Figures and table for "Microbiota-derived indole limits *Campylobacter jejuni* colonization by inhibiting respiration and metabolism": Supplementary Table 1.pdf

**Supplementary Table 1: Strains, Plasmid and Primers:**

| **Strain or Plasmid** | **Relevant Characteristics** | **Source or reference** |
|---|---|---|
| *C. jejuni* NCTC 11168 (WT) | Wild type strain from human gastroenteritis stool | |
| *C. jejuni* 11168 l*ctP*::kan | *lctP* insertion mutant of WT 11168 strain (KanR) | (3) |
| *C.jejuni* 11168 n*apA*::cm | *napA* insertion mutant of WT 11168 strain (cmR) | This study |
| *C.jejuni* 11168 *ackA::pta::*cm | ackA insertion mutant of WT 11168 strain (cmR) | This study |
| *C.jejuni 11168 motAB::kan* | motAB insertion mutant of WT 11168 strain (kanR) | Kind gift from Prof **Christine M. Szymanski** |
| DRH212 | Streptomycin resistant of Wild type *C. jejuni* strain 81176 | |
| *C.jejuni* ATCC 43431 | Wild type strain isolated from clinical stool samples | |
| *C. jejuni* TW 19241 | Wild type multidrug resistant strain from stool sample | Kind gift from Prof. Shannon Manning (MSU). |
| *E.coli* Nissle 1917 (EcN) | Wild type strain Probiotic strain streptomycin resistant | Kind gift from Prof. Jeffery H Withey (Wayne state University) |
| *E.coli* Nissle 1917 (EcN) pKD46 | EcN strain carrying red recommibanse plasmid pKD46 (ampR) | This study |
| *E.coli* Nissle 1917 (EcN) tnaA::cm | tnaA insertion mutant of EcN (cmR) | This study |
| pGEMT | pGEMT Subcloning vector (AmpR) | |
| pRY109 | Contains Campylobacter Chloramphenicol Crassest | |
| pKD46 | the expression of Red recombinase | |
| pKD4 | obtain the chloramphenicol acetyl transferase gene (*cat*) Chloramphenicol resistant gene | |

| Primers | Sequence |
|---|---|
| NapA knock out Primer A: | 5'-CCG CTA TTG CAA GTG CTG CTA GTG -3' |
| NapA knock out Primer B: | 5'-CCCATCCACTATAAACTAACA CCCGGGGGAC CAA AGG ATT GGG TGC A -3' |
| NapA knock out Primer C: | 5'-TGTTAGTTTATAGTGGATGGGTCCCGGGTGTC TTA CCT GTA GGT GCT GCC -3' |
| NapA knock out m Primer D: | 5'- GGC ACA CGC ATA GTC ATA GTT CC -3' |
| acka knock out primer A: | 5'- AGG CAT GAG TTT AGA AGA GGC TAA GAT GC -3' |
| ackA knock out primer B: | CCCATCCACTATAAACTAACA CCCGGGG CAA TAT CAT CTA CCA AAC ATC CAC GAC |
| ackA knock out Primer C: | TGTTAGTTTATAGTGGATGGGCCCGGGAAT GCG CTT GAT GGC TGC GGT CAT CGT |
| ackA knock out Primer D: | GCT TTA TCA TTA CCT TGT TCT ATC TCT C |
| TnaA Knock out F: | 5'- TGA AGA GGC AAT TAT TAA ATC CGG CAT GAA CCC GTT CCT GC GTG TAGGCT GGA GCT GCT TC -3' |
| TnaR Knock out R: | 5'- AGT GAC GCA ATA CTT TCG GTT CGT AGG TAA AGG TTA ACC CCA TAT GAA TAT CCT CCT TAG |
| **Real Time PCR Primers** | |
| napA F | 5'-GAAAGCACCTGAAAGACCAAG-3' |
| napA R | 5'-ACACGCATAGTCATAGTTCCAC-3' |
| ccoN F: | 5'-GACTTAGACCACTTCATACTTCAGG-3' |
| ccoN R: | 5'-CCATACTCACTTTAAGAACACGC-3' |
| RpoA F: | 5'-GAGATCGGCTATGGAATCACC-3' |
| RpoA R: | 5'-CATCTTCAAGCATACCACGC-3' |
| lctP F: | 5'-TGCTGTTCTATGCCGACAAG-3' |
| lctP R: | 5'- TCTTATCGCAAATACCGCTCC-3' |
| Pta F | 5'-GCTGATGCTATGGTTAGTGGAG-3' |
| Pta R | 5'-TCCTGAAACGGAACTAATACCTTC-3' |
| AckA F | 5'-CACCTAGTGTTGCAAATGTGG-3' |
| AckA R | 5'-GACAAAAGCATGAGAAGTCCC-3' |
| **Mouse Real time PCR Primers** | |

| | |
|---|---|
| *Nos2 F:* | CCTCTTTCAGGTCACTTTGGTAGG |
| Nos2 R | TTGGGTCTTGTTCACTCCACGG |
| KC F: | 5’- GCTTGCCTTGACCCTGAAGCTC-3’ |
| KC R | 5’- TGTTGTCAGAAGCCAGCGTTCAC-3’ |
| Tnf alpha F: | 5'-TCGAGTGACAAGCCTGTAGCC-3' |
| Tnf alpha R: | 5'-TTGAGATCCATGCCGTTGG-3' |
| IL-6 F: | 5'-GGATACCACTCCCAACAGACC-3' |
| mIL-6 R: | 5'-GTGCATCATCGTTGTTCATACAA-3' |
| Actin F | CTTCTTTGCAGCTCCTTCGTT |
| Actin R | AGGAGTCCTTCTGACCCATTC |
| Ahr F | GGCTTTCAGCAGTCTGATGTC |
| Ahr R | CATGAAAGAAGCGTTCTCTGG |
| IL-10 F | AACTGCACCCACTTCCCAGTC |
| IL-10 R | CATTAAGGAGTCGGTTAGCAG |
