## Supplementary Figures and table for "Microbiota-derived indole limits *Campylobacter jejuni* colonization by inhibiting respiration and metabolism": Supplementary Table 2.pdf

| Down regulated genes in presence of Indole | | |
|---|---|---|
| **Gene names/ Function** | **Fold changes (log 2)** | ***P* Value** |
| **Respiration and metabolism** | | |
| **Nitrate reductase** | | |
| *napB* | -2.7810548 | *** |
| *napG* | -2.7422321 | *** |
| *napA* | -2.4305909 | *** |
| *napL* | -1.5500835 | *** |
| **Nitrite reductase** | | |
| *nrfH* | -4.9617015 | *** |
| *nrfA* | -4.3584494 | *** |
| **Cbb3- Cytochrome C terminal oxidase (aerobic respiration)** | | |
| *ccoN* | -1.8336232 | *** |
| *ccoP* | -1.5496148 | *** |
| *ccoO* | -2.3831166 | *** |
| *ccoQ* | -1.9677355 | *** |
| **Hydrogenase** | | |
| *hydB* | -1.1877599 | *** |
| *hydA* | -1.1816983 | *** |
| **Format dehydrogenase** | | |
| *fdhA* | -1.7391597 | *** |
| *fdhB* | -1.3672822 | *** |
| **TCA cycle regulatory genes** | | |
| *gltA* | -2.6730476 | *** |
| *acnB* | -2.3307891 | *** |
| *oorC* | -1.1784371 | *** |
| *fumC* | -1.1312133 | *** |
| *mqo* | -1.472393 | *** |
| *sucC* | -1.152045 | *** |
| *pycA* | -1.2666124 | *** |

| | | |
|---|---|---|
| *frdC* | -1.2867861 | *** |
| *frdB* | -2.2634438 | *** |
| *frdA* | -2.4802501 | *** |
| *aspA* | -3.5216678 | *** |
| *aspB* | -2.0188467 | *** |
| **Acetate switch (Acetyl-coA to Acetate)** | | |
| *pta* | -1.98407 | *** |
| *ackA* | -1.4124585 | *** |
| **Lactate transporter and metabolism pathways** | | |
| *lctP* | -1.9105147 | *** |
| *Cj0075c* | -2.3093887 | *** |
| *Cj0074c* | -2.3202608 | *** |
| *Cj0073c* | -2.9362736 | *** |
