## Supplementary Figures and table for "Microbiota-derived indole limits *Campylobacter jejuni* colonization by inhibiting respiration and metabolism": Supplementary Table 3 Up regulated genes in presence of Indole.pdf

| Up regulated genes in presence of Indole | | |
|---|---|---|
| **Gene names/ Function** | **Fold changes (log 2)** | ***P* Value** |
| **Oxidative stress/ Acid stress** | | |
| *katA* | + 4.56693691 | *** |
| *sodB* | +1.92135053 | *** |
| *ahpC* | +2.391672773 | *** |
| *perR* | +1.86655541 | *** |
| *Cj1659 (p19)* | +4.08544019 | *** |
| **Putative ABC-transporter** | | |
| *Cj1663* | +2.890346073 | *** |
| *Cj1662* | +4.900037704. | *** |
| *Cj1661* | +4.613569626 | *** |
| *Cj1660* | +4.792500289 | *** |
| **Phosphate Uptake & Nutrient Transport** | | |
| *pstS* | +3.791743131 | *** |
| *pstC* | +4.755112856 | *** |
| *pstA* | +3.089200668 | *** |
| **Gluconate dehydrogenase** | | |
| *Cj0414* | +3.030565429 | *** |
| *Cj0415* | +3.565323375 | *** |
