## Supplementary figures and images for "Microbiota-derived indole limits *Campylobacter jejuni* colonization by inhibiting respiration and metabolism"

### Sfig2.jpg.pdf

#### Supplementary Figure 2:

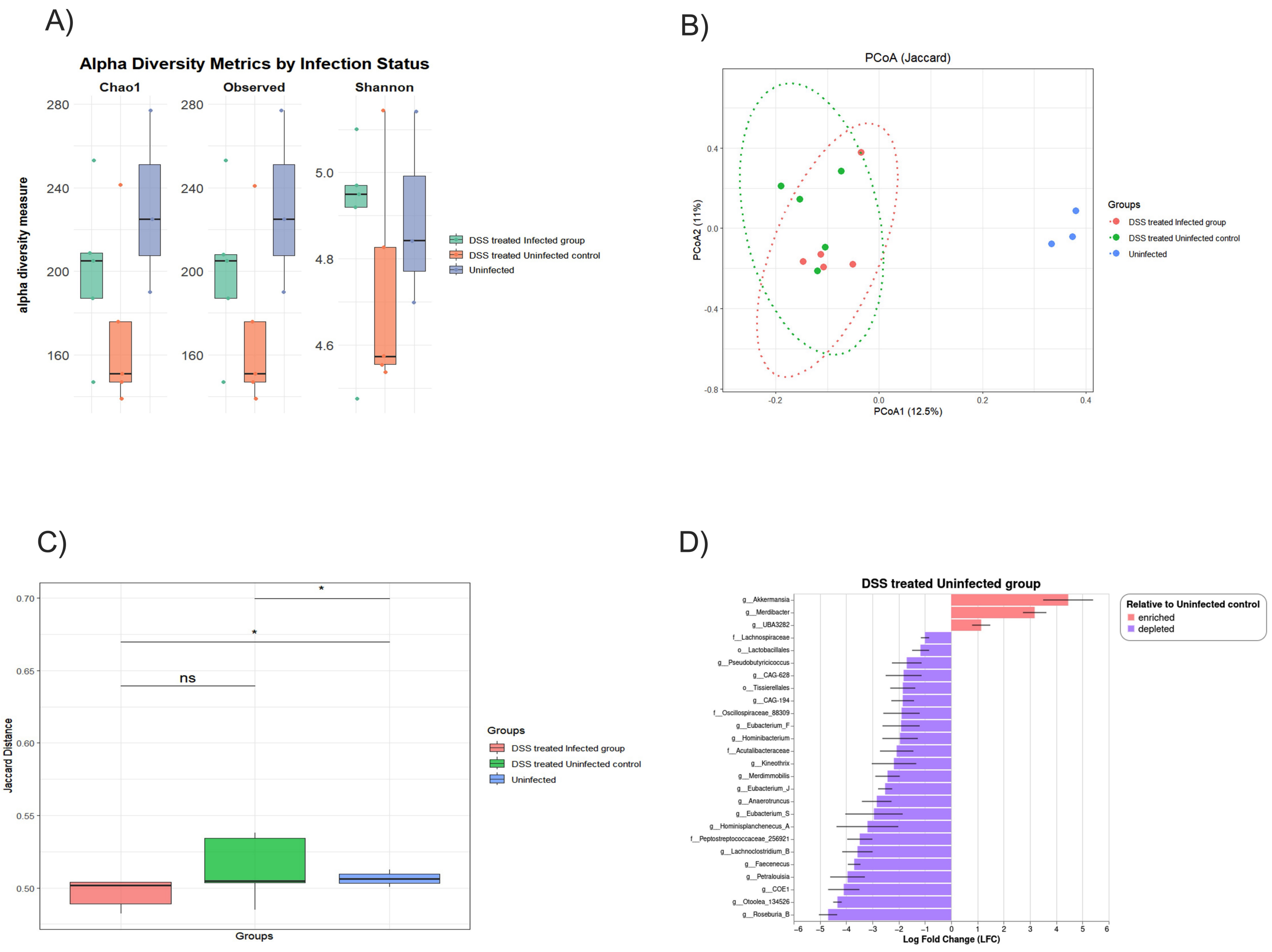

### sFig3.jpg.pdf

#### Supplementary Fig 3

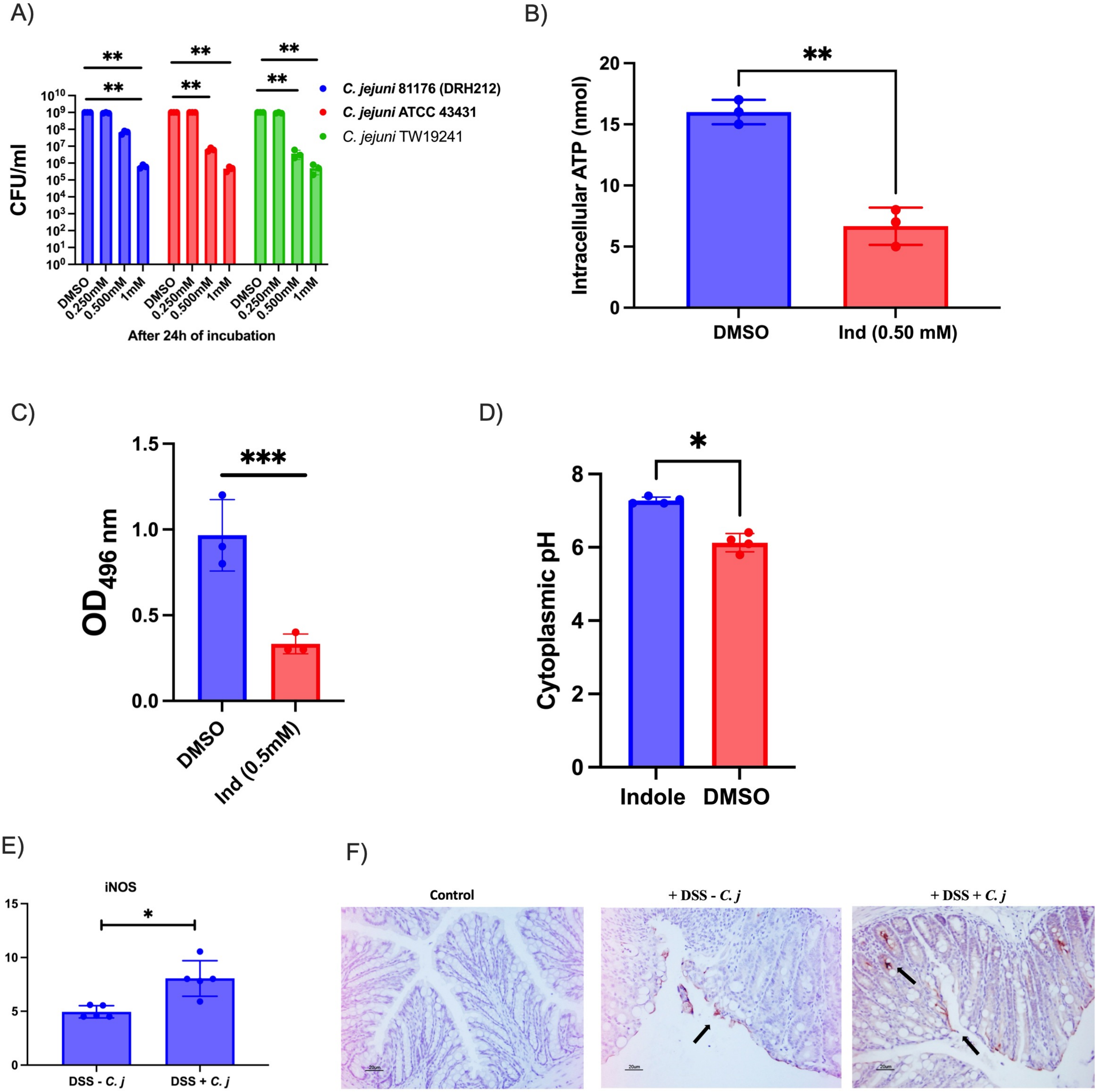

### sfig 1.jpg.pdf

#### Supplementary Fig 1:

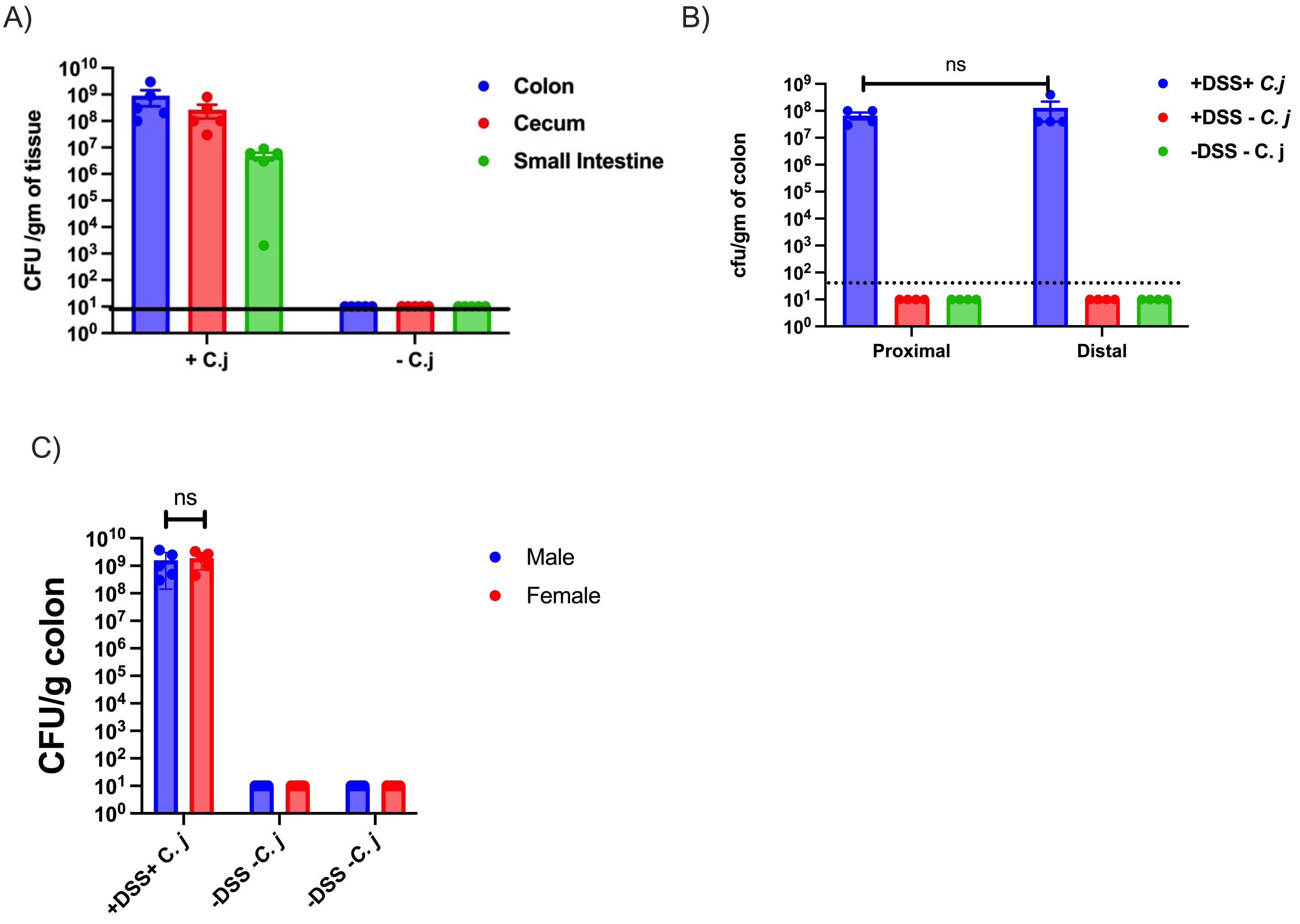
